# BRD4 Interacts with GATA4 to Govern Mitochondrial Homeostasis in Adult Cardiomyocytes

**DOI:** 10.1101/2020.04.16.041806

**Authors:** Arun Padmanabhan, Michael Alexanian, Ricardo Linares-Saldana, Bárbara González-Terán, Gaia Andreoletti, Yu Huang, Andrew J. Connolly, Wonho Kim, Austin Hsu, Qiming Duan, Sarah A. B. Winchester, Saptarsi M. Haldar, Rajan Jain, Deepak Srivastava

**Author notes:** These authors contributed equally to this work (Arun Padmanabhan, Michael Alexanian). Address correspondence to: Deepak Srivastava, Gladstone Institute of Cardiovascular Disease, 1650 Owens St, San Francisco, CA 94158, Rajan Jain, Perelman School of Medicine, 3400 Civic Center Blvd, Philadelphia, PA 19104.

## Abstract

Gene regulatory networks control tissue plasticity during basal homeostasis and disease in a cell-type specific manner. Ubiquitously expressed chromatin regulators modulate these networks, yet the mechanisms governing how tissue-specificity of their function is achieved are poorly understood. BRD4, a member of the BET (Bromo- and Extra-Terminal domain) family of ubiquitously expressed acetyl-lysine reader proteins, plays a pivotal role as a coactivator of enhancer signaling across diverse tissue types in both health and disease, and has been implicated as a pharmacologic target in heart failure. However, the cell-specific role of BRD4 in adult cardiomyocytes remains unknown. Here, we show that cardiomyocyte-specific deletion of BRD4 in adult mice leads to acute deterioration of cardiac contractile function with mutant animals demonstrating a transcriptomic signature enriched for decreased expression of genes critical for mitochondrial energy production. Genome-wide occupancy data show that BRD4 enriches at many downregulated genes and preferentially co-localizes with GATA4, a lineage determining cardiac transcription factor not previously implicated in regulation of adult cardiac metabolism. Co-immunoprecipitation assays demonstrate that BRD4 and GATA4 form a complex in a bromodomain-independent manner, revealing a new interaction partner for BRD4 that has functional consequences for target transactivation and may allow for locus and tissue specificity. These results highlight a novel role for a BRD4-GATA4 module in cooperative regulation of a cardiomyocyte specific gene program governing bioenergetic homeostasis in the adult heart.

## Introduction

Heart failure (HF) is a clinical syndrome that occurs when a weakened heart is unable to maintain organ perfusion at a level adequate to meet tissue demand, resulting in shortness of breath, fatigue, and early death. This condition represents an enormous burden to the healthcare system accounting for almost 2% of all medical expenditures annually and, despite the current standard of care, carries a dismal prognosis with a 5-year mortality approaching 50%^1,2^. The mainstays of currently approved pharmacologic therapy for HF target neurohormonal signaling pathways with beta-adrenergic receptor antagonism, inhibition of the renin-angiotensin system, and augmentation of the natriuretic peptide system, all of which have improved survival in HF patients^3^. Despite these successes, the residual burden of morbidity and mortality in HF remains unacceptably high, underscoring an urgent need for novel treatment approaches^1^.

During HF pathogenesis, hemodynamic and neurohormonal stressors activate a network of signal transduction cascades that ultimately converge upon the nucleus where, specific transcription factors (TFs) function in the context of chromatin to drive maladaptive gene expression programs and modulate cell state^4,5^. In response, the heart undergoes a stereotypical pathologic remodeling process characterized by cardiomyocyte (CM) hypertrophy, interstitial fibrosis, and altered substrate/energy utilization that culminates in organ-level contractile dysfunction. Studies in animal models of HF have implicated several nodal TFs (e.g., NFAT, GATA4, MEF2, and NF-κB) as drivers of disease progression through their induction of gene expression programs that may provide short-term adaptation to pathologic stress but whose sustained activation progressively weakens cardiac performance^4,6^. This stress-coupled activation of DNA binding proteins in HF elicits global changes in chromatin structure and post-translational modifications on histone proteins, including lysine acetylation of histone tails, TFs, and other chromatin associated proteins. Given the central role of signal-coupled gene transcription in cardiac plasticity, manipulation of chromatin-dependent signaling as a therapeutic approach for HF has been an area of intense interest^7,8^.

Previous work has established a crucial role for the BET (Bromo- and Extra-Terminal domain) family of acetyl-lysine “reader” proteins in the epigenetic control of adverse cardiac remodeling and HF pathogenesis^9,10^. There are four mammalian BET proteins (BRD2, BRD3, and BRD4 that are ubiquitously expressed and BRDT, which is testis-specific) that each contain two tandem N-terminal bromodomains (BDs) that mediate acetyl-lysine binding^11,12^. BRD4 is the most studied member of this family and is a highly pursued target in cancer^13,14^. BRD4 associates with acetylated chromatin at active enhancers and promoters, where it interacts with the transcriptional machinery to signal to RNA polymerase II and activate transcription^15^. Within the BET family, BRD4 is unique in that it contains an extended C-terminal domain (CTD) that interacts with the cyclin T1 and CDK9 subunits of the positive transcription elongation factor b (PTEFb) complex and functions as an allosteric coactivator of transcription elongation^16,17^. The ability to probe BET function in mammalian biology was rapidly accelerated by the creation of JQ1, a potent and specific small molecule tool compound that reversibly binds the dual BDs of all BET proteins with high affinity via exquisite shape complementarity to the acetyl-lysine binding pocket of these proteins^18,19^. JQ1 competitively and reversibly displaces BET proteins from their acetyl-lysine interaction partners on active enhancers (e.g., acetylated histones or acetylated TFs), thereby disrupting signaling between enhancers and promoters^18,19^. Prior studies have demonstrated that systemic delivery of JQ1 potently ameliorates hallmark features of HF pathogenesis in an array of rodent models, including those mediated by pressure overload, adrenergic agonists, myocardial infarction, and dilated cardiomyopathy (DCM)-causing variants^9,10,20,21^. However, the precise identity of cell-types and BET isoforms that mediate these therapeutic benefits remain a major unanswered question with important translational implications. As systemic delivery of pan-BET inhibitors such as JQ1 are unable to probe the gene-specific and cell-compartment-specific functions of BRD4 *in vivo*, we set out to discover the role of BRD4 in adult CMs using a conditional genetic approach in mice.

Here, we identify a critical role for BRD4 in maintaining murine cardiac homeostasis *in vivo*. As germline deletion of *Brd4* results in early zygotic implantation defects and haploinsufficiency results in severe multisystem developmental abnormalities^22^, we generated a novel *Brd4* conditional allele and genetically deleted *Brd4* in adult murine CMs. Tamoxifen-inducible genetic ablation of *Brd4* in these cells resulted in a rapid and severe reduction in global left ventricular (LV) systolic function, LV cavity dilation, and uniform death. Integration of gene expression, genomic occupancy, and chromatin accessibility datasets demonstrated that BRD4 regulates mitochondrial gene expression. These analyses unveiled a new functional interaction between BRD4 and GATA4 in regulating mitochondrial gene expression and CM homeostasis. We confirmed a physical interaction between BRD4 and GATA4 that occurs in a BD-independent fashion. Taken together, our data reveal that BRD4 is a critical regulator of basal CM homeostasis and identify a potential direct role for GATA4 in the regulation of metabolic gene programs in the adult heart. In addition, these findings highlight key differences between gene deletion and chemical biological approaches using compounds such as JQ1.

## Results

### Loss of BRD4 in adult CMs results in contractile dysfunction and lethality

In order to decipher tissue-specific roles for BRD4 *in vivo*, we generated a new *Brd4* conditional allele by engineering a homologous recombination event in mouse embryonic stem (ES) cells (**Figure S1A**) in a pure C57BL/6 background. We designed a targeting strategy that inserted flanking loxP sites around the third exon of *Brd4*, which contains the canonical translational start site, with loxP sites and verified appropriate targeting in several ES cell clones by Southern blotting (**Figure S1B,C**). These were used to generate chimeric mice that were subsequently bred for germline transmission. Mice harboring this allele (designated *Brd4^flox^*) were normal, viable, and able to breed in both the heterozygous and homozygous state (**Figure S1D**). These mice were crossed to mice harboring the Myh6-Mer-Cre-Mer (*Myh6-MCM*) allele^23^, which permits for tamoxifen (TAM)-inducible CM-specific deletion of *Brd4*. Immunofluorescence of heart tissue (**Figure S2**) and immunoblotting of isolated CM lysates (**Figure 1A**) confirmed efficient loss of BRD4 protein abundance after 5 days of TAM administration.

**Figure 1:**
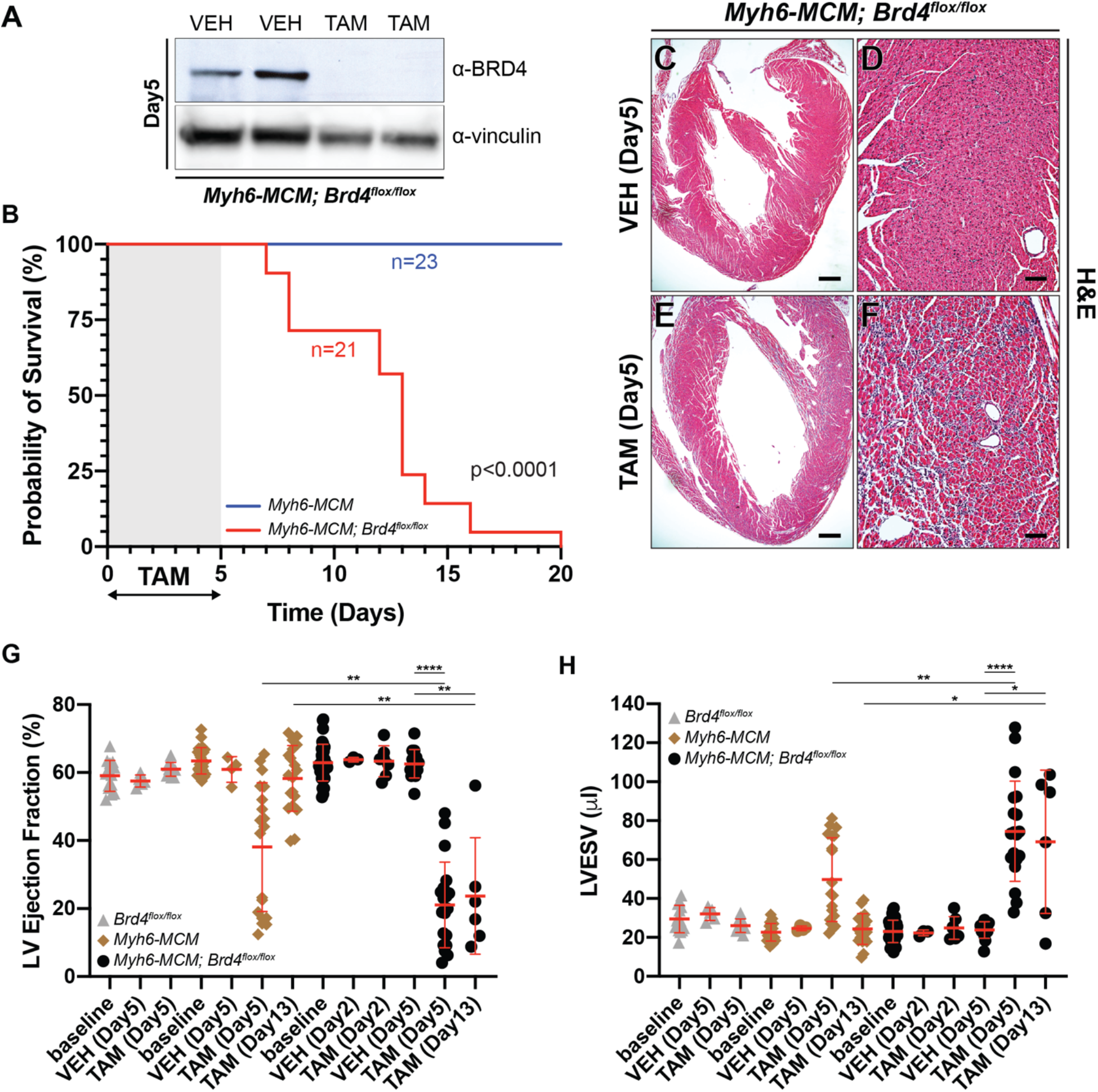
Adult cardiomyocyte-specific *Brd4* deletion results in acute and persistent contractile dysfunction and lethality. **(A)** Immunoblot of isolated *Myh6-MCM; Brd4^flox/flox^* cardiomyocyte lysates from mice treated with vehicle (VEH) or tamoxifen (TAM) for 5 days using BRD4 or vinculin (loading control) antibodies. **(B)** Kaplan-Meier curve demonstrating survival of indicated mice treated with tamoxifen (TAM; 75 μg/g/day). **(C-F)** H&E images of *Myh6-MCM; Brd4^flox/flox^* mice treated with vehicle (VEH) or tamoxifen (TAM) at low and high magnification. **(G)** Ejection fraction and **(H)** left ventricular end systolic volume (LVESV) of indicated mice treated with tamoxifen (TAM) or vehicle (VEH) at indicated days after injection. Individual points and mean ± 1 SD shown. ** indicates p<0.0027, **** indicates p<0.00001. Scale bars = 100 μm (D, F) and 500 μm (C, E).

Following administration of intraperitoneal TAM (75 μg/g/day) or vehicle (VEH; corn oil) for 5 consecutive days to an initial cohort of adult mice, we noticed that *Myh6-MCM; Brd4^flox/flox^* mice became lethargic and ill-appearing, and several succumbed shortly thereafter. In a survival analysis of *Myh6-MCM, Brd4^flox/flox^* (n=21) animals treated with TAM, 83% died within 14 days of the last dose of TAM with 100% mortality by 21 days. In contrast, 100% of TAM-treated *Myh6-MCM* (n=23) animals survived (**Figure 1B**). Histological analyses revealed interstitial infiltrates in TAM treated *Myh6-MCM; Brd4^flox/flox^* animals when compared with VEH-treated controls of the same genotype (**Figure 1E,F vs. C,D**), but we did not find any significant differences in apoptosis (as measured by cleaved caspase-3 immunofluorescence staining) when measured at day 5 post-TAM initiation (**Figure S3**).

We next assessed cardiac function by transthoracic echocardiography. We generated cohorts of *Brd4^flox/flox^* (n=14), *Myh6-MCM* (n=23), or *Myh6-MCM; Brd4^flox/flox^* mice (n=41) at 8-12 weeks of age, which were then administered intraperitoneal TAM (75 μg/g/day) or VEH (corn oil) for 5 consecutive days. Echocardiography was performed at baseline, and on day 2, 5, and 13 after commencing TAM. CM-specific deletion of *Brd4* resulted in a significant reduction in LV ejection fraction (EF) when compared with VEH (21% vs. 63%, p<0.0001) and LV chamber dilation (LV end systolic volume [LVESV] 24 μl vs. 75 μl, p<0.0001) (**Figures 1G,H**) after 5 days of TAM administration. No appreciable changes in these indices were detected following 2-days of TAM treatment (EF 63% vs 63%, ns; LVESV 22 μl vs. 25 μl, ns). Treatment of *Brd4^flox/flox^* animals with TAM or VEH did not result in any significant changes in these echocardiographic parameters. Consistent with previous reports^24^, administration of TAM alone for 5 days in *Myh6-MCM* animals led to a transient decrease in LV systolic function (EF 38% vs. 61%, p<0.0001) and LV chamber dilation immediately following TAM exposure (LVESV 25 μl vs. 50 μl, p<0.0001). However, the degree of LV systolic dysfunction and chamber dilation was significantly greater in *Myh6-MCM; Brd4^flox/flox^* animals when compared to *Myh6-MCM* controls at the 5 day time point (EF 21% vs. 38%, p<0.0027; LVESV 75 μl vs. 50 μl, p<0.0028). Importantly, repeat echocardiographic analysis at day 13 revealed normalization of these indices in *Myh6-MCM* controls consistent with a transient and reversible Cre-mediated toxicity. In contrast, *Myh6-MCM; Brd4^flox/flox^* animals that survived to this time point failed to demonstrate any recovery (**Figure 1G,H**). These echocardiographic data are consistent with our finding that only the TAM-treated *Myh6-MCM; Brd4^flox/flox^* mice, and not the equivalently treated *Myh6-MCM* mice, had a striking propensity for early mortality after TAM treatment (**Figure 1A**). Taken together, we demonstrate that postnatal deletion of *Brd4* in CMs of adult mice leads to early mortality and severe systolic HF within 5 days that is sustained, suggesting an essential role for BRD4 in maintaining expression of key homeostatic gene programs in the adult CM.

### BRD4 regulates mitochondrial bioenergetic gene pathways

To better understand the mechanism by which BRD4, a potent transcriptional coactivator, regulates homeostasis in adult CMs, we performed bulk RNA-sequencing (RNA-seq) on isolated CMs from *Myh6-MCM; Brd4^flox/flox^* treated with TAM (*Brd4*-KO) or VEH (control) and *Myh6-MCM* mice treated with TAM (Cre-control) (**Figure S4A,B**). We collected samples at two early time points that occurred prior to the onset of mortality (n≥2 biological replicates per condition): day 2 post-TAM (prior to the decrease in LVEF) and day 5 post-TAM (when acute HF was first detected).

While Cre-control mice did not display mortality, we noted transient decrease in LV systolic function in these mice after 5 days of TAM treatment. Therefore, we sought to rigorously and quantitatively correct for any degree of gene dysregulation that was specifically associated with transient Cre-activation. Using a permissive statistical threshold to liberally capture the vast majority of Cre-related effects (Log2 FC±1, adjusted p<0.05), we found that Cre-control CMs demonstrated differential expression of 5,745 genes compared with control CMs at the 5 day time point (**Figure S4C,D**). Gene ontology (GO) analysis of those genes downregulated in Cre-control CMs were enriched for terms relevant to regulation of CM contractile activity and energy handling whereas those associated with upregulated genes were enriched for programs that control ribosome function (**Figure S4D**). The transient myocardial dysfunction specifically associated with Cre-activation is consistent with prior reports^24^ and we highlight the transcriptomic effects of this manipulation, which may serve as a resource for the field to adjust RNA-seq signatures for the potentially confounding effects of Cre-activation in adult CMs. Using these data, we defined a molecular signature following 2- and 5-days of tamoxifen treatment in *Myh6-MCM* CMs and quantitatively subtracted this from all subsequent analyses comparing *Brd4*-KO and control CMs (**Table S1-4**).

Comparison of *Brd4*-KO and control samples at day 2 and 5 reveal a temporal increase in the number of dysregulated genes, consistent with the degree of cardiac dysfunction at each time point. We observed 595 dysregulated genes at day 2, prior to cardiac dysfunction. GO analysis of the day 2 differentially expressed (DE) gene set yielded heterogeneous terms with significant p-values, with downregulation of genes involved in cardiac contraction and cardiac development (**Figure 2A**). By day 5, *Brd4*-KO CMs demonstrated differential expression of 2,157 genes when compared with controls (807 upregulated, 1,350 downregulated, Log2 FC±1, p<0.05) (**Figure 2B**), after stringent subtraction of Cre-only dysregulation. Importantly, GO analysis revealed a striking signature for mitochondrial bioenergetics amongst those genes preferentially downregulated in *Brd4*-KO CMs, a pathway wherein BRD4 has not been explicitly implicated. GO analysis of those genes upregulated in *Brd4*-KO CMs were enriched broadly for general terms associated with cardiac stress, including fibrotic and inflammatory cellular processes, suggesting that many of these upregulated genes may be a secondary response to acute loss of BRD4 coactivator function in CMs (**Figure 2B**).

**Figure 2:**
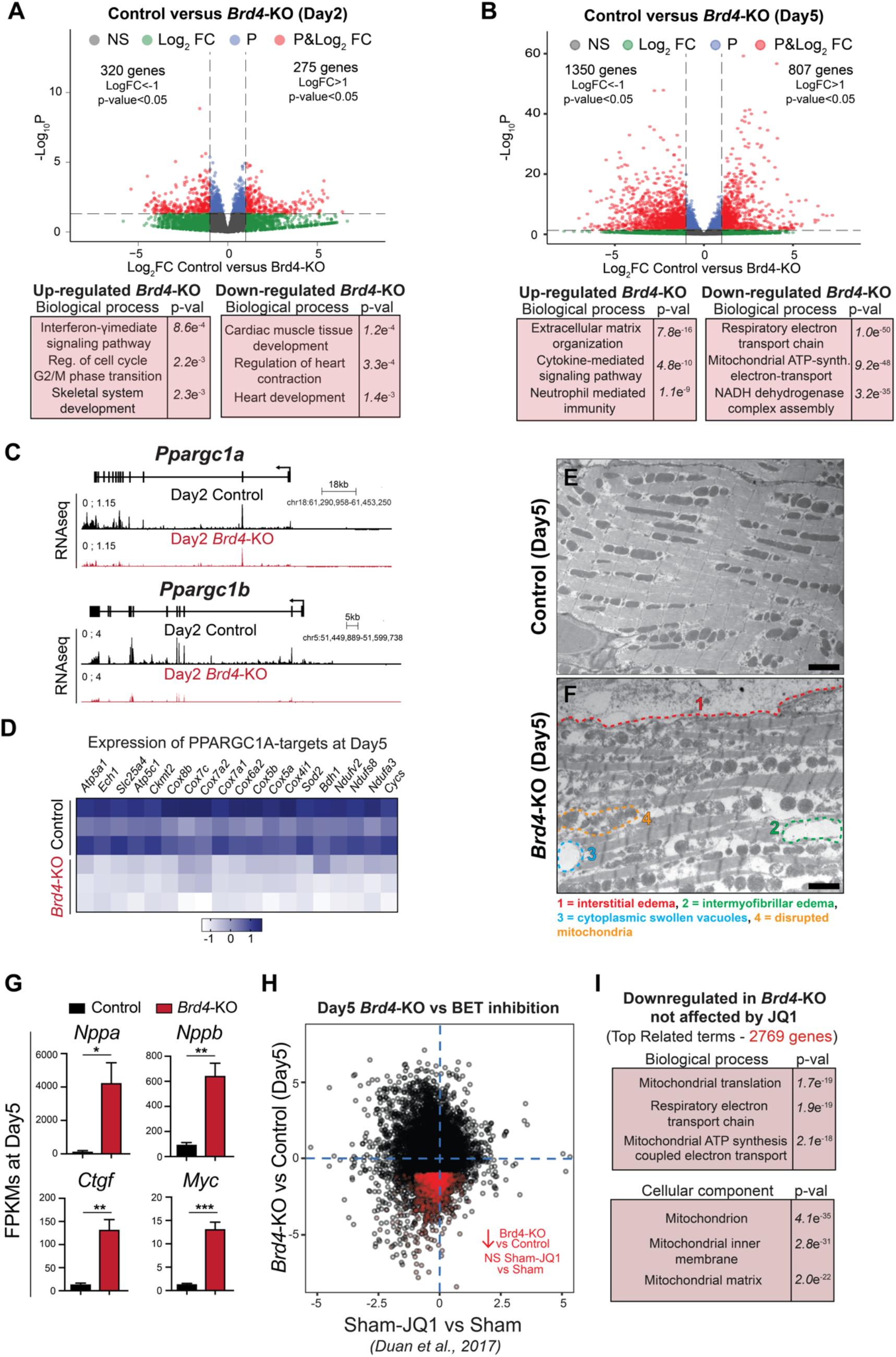
BRD4 regulates mitochondrial metabolic pathways in adult cardiomyocytes. **(A, B)** Volcano plots showing fold change and Log2 p-value of individual genes 2 days **(A)** or 5 days **(B)** after *Brd4* deletion; genes differentially expressed between Cre-control and Control samples have been excluded. Selected categories identified from gene ontology analysis from up or down regulated genes are shown below the volcano plot. **(C)** Track view of *Ppargc1a* and *Ppargc1b* genes showing sequencing reads mapping from RNAseq signature at day 2 post-tamoxifen (TAM) treatment for Control and *Brd4*-KO samples. **(D)** Heatmap of expression of PPARGC1A known targets^25^ in Control and *Brd4*-KO samples at day 5 post-TAM treatment. **(E, F)** Electron micrographs of *Brd4*-KO and Control animals at day 5 highlighting the loss of normal mitochondrial morphology. **(G)** FPKMs of indicated genes related to cardiac stress and homeostasis in Cre-control, Control, or *Brd4*-KO cardiomyocytes at day 5. Error bars represent standard error of the mean (SEM) (* indicates p<0.05; ** indicates p<0.01). **(H)** Correlation analysis of difference in gene expression in day 5 *Brd4*-KO cardiomyocytes compared to sham-operated animals administered JQ1 (normalized to their respective controls). Genes highlighted in red are those which are downregulated upon BRD4 loss but not significantly changed by JQ1. **(I)** Gene ontology analysis of genes that were downregulated upon BRD4 loss but not affected by JQ1 treatment. Scale bars = 2 μm (E, F).

Given the marked transcriptional dysregulation of gene programs linked to mitochondrial homeostasis in *Brd4*-KO CMs at day 5, we turned our attention to the earliest time point in our analysis (day 2) to interrogate if established nodal regulators of mitochondrial metabolism were downregulated following *Brd4* loss. Indeed, we discovered that the expression of the transcriptional co-activators *Ppargc1a* (PPARGC1A; PGC-1α) and *Ppargc1b* (PPARGC1B; PGC-1β) were significantly downregulated in *Brd4*-KO CMs when compared to control CMs at this early time point, prior to the appearance of any LV systolic dysfunction (**Figure 2C**). PGC-1α/β are known to be master regulators of mitochondrial biogenesis and oxidative phosphorylation gene programs in both cardiac and skeletal muscle cells^34,35^, suggesting the mitochondrial dysregulation may be a primary consequence of *Brd4* deletion. At day 5, expression of known targets of PGC-1α in the heart^34^ were downregulated in *Brd4*-KO CMs when compared with controls, including genes involved in oxidative phosphorylation, fatty acid oxidation, and ATP synthesis (**Figure 2D**). Given this signature of broad mitochondrial dysfunction identified at the transcriptomic level, we performed electron microscopy to assess cardiac ultrastructure and mitochondrial morphology following BRD4 loss. While control samples showed an organized arrangement of myofibers with normal sarcomeres (**Figure 2E**), *Brd4*-KO hearts demonstrated interstitial edema between CMs (**Figure 2F,1**), intermyofibrillar edema within CMs (**Figure 2F,2**), swollen cytoplasmic vacuoles (**Figure 2F,3**), and disrupted mitochondria that showed mild swelling (**Figure 2F,4; Figure S5**). These findings are often observed with severe disruption of cellular metabolism, underscoring a CM-intrinsic gene program controlling basal mitochondrial energetics in the adult heart that is acutely sensitive to BRD4 abundance.

Recent reports demonstrate that pharmacologic BET BD inhibition with JQ1 ameliorates adverse cardiac remodeling and HF in several murine models with JQ1-treated animals showing improved function and decreased inflammation and fibrosis^9,10,20^. However, JQ1-treatment did not affect exercise-induced physiological cardiac hypertrophy^20^, a form of plasticity which features upregulation of genes involved in mitochondrial biogenesis and oxidative metabolism. Transcriptional profiling of the sham-operated animals treated with JQ1 did not reveal dramatic changes, suggesting that BET BD inhibition in homeostatic conditions is not associated with a pronounced gene dysregulation signature in the heart^20^. This is in stark contrast to the broad transcriptional changes that follow *Brd4* CM deletion that results in marked upregulation of canonical cardiac stress markers, including *Nppa*, *Nppb*, *Ctgf,* and *Myc* (**Figure 2G**) with concomitant downregulation of mitochondrial and metabolic genes. To assess this quantitatively, we compared gene expression changes seen with JQ1-treatment in sham-operated mice to those that follow CM-specific loss of *Brd4* in adult mice. We confirmed that global gene expression changes were poorly correlated between pharmacologic BET BD inhibition and BRD4 loss at day 5 post-TAM administration (**Figure 2H**). GO analysis of those genes downregulated in *Brd4-*KO CMs whose expression was unaffected by JQ1 treatment were enriched specifically for mitochondrial-related terms (**Figure 2I**), highlighting that transient exposure to small molecule BET BD inhibitors such as JQ1 and *Brd4* CM-deletion are two very different molecular perturbations.

### BRD4 colocalizes with GATA4 at genes controlling mitochondrial bioenergy production

Given the specific changes in gene expression caused by BRD4 deletion and the known role of BRD4 as a coactivator of transcription, we queried changes in chromatin accessibility following CM-specific BRD4 loss. We performed Assay for Transposase-Accessible Chromatin followed by sequencing (ATAC-seq)^25^ on either control or *Brd4*-KO CMs isolated at day 5 (n=2 replicates per condition). We found that 58% of ATAC-seq peaks identified in control and *Brd4*-KO samples were shared (13,428 of 23,052 total peaks) (**Figure 3A**). In addition, we found a substantial number of accessible regions unique to *Brd4*-KO samples (n=5,242 peaks) or control samples (n=4,328 peaks), defined as regions of the genome which became accessible (gained in *Brd4*-KOs) or lost accessibility (lost in *Brd4*-KOs) upon BRD4 loss, respectively (**Figure 3A**). Ontology analysis of the regions which lose accessibility in *Brd4*-KO CMs showed enrichment for regulatory elements of genes linked to CM identity (e.g., sarcomere organization and myofibril assembly) and mitochondrial function, suggesting that BRD4 is required for those regions to be accessible (**Figure 3B**). Interestingly, accessible regions gained in *Brd4*-KO CMs were related to elements associated with stress responses (**Figure 3B**).

**Figure 3:**
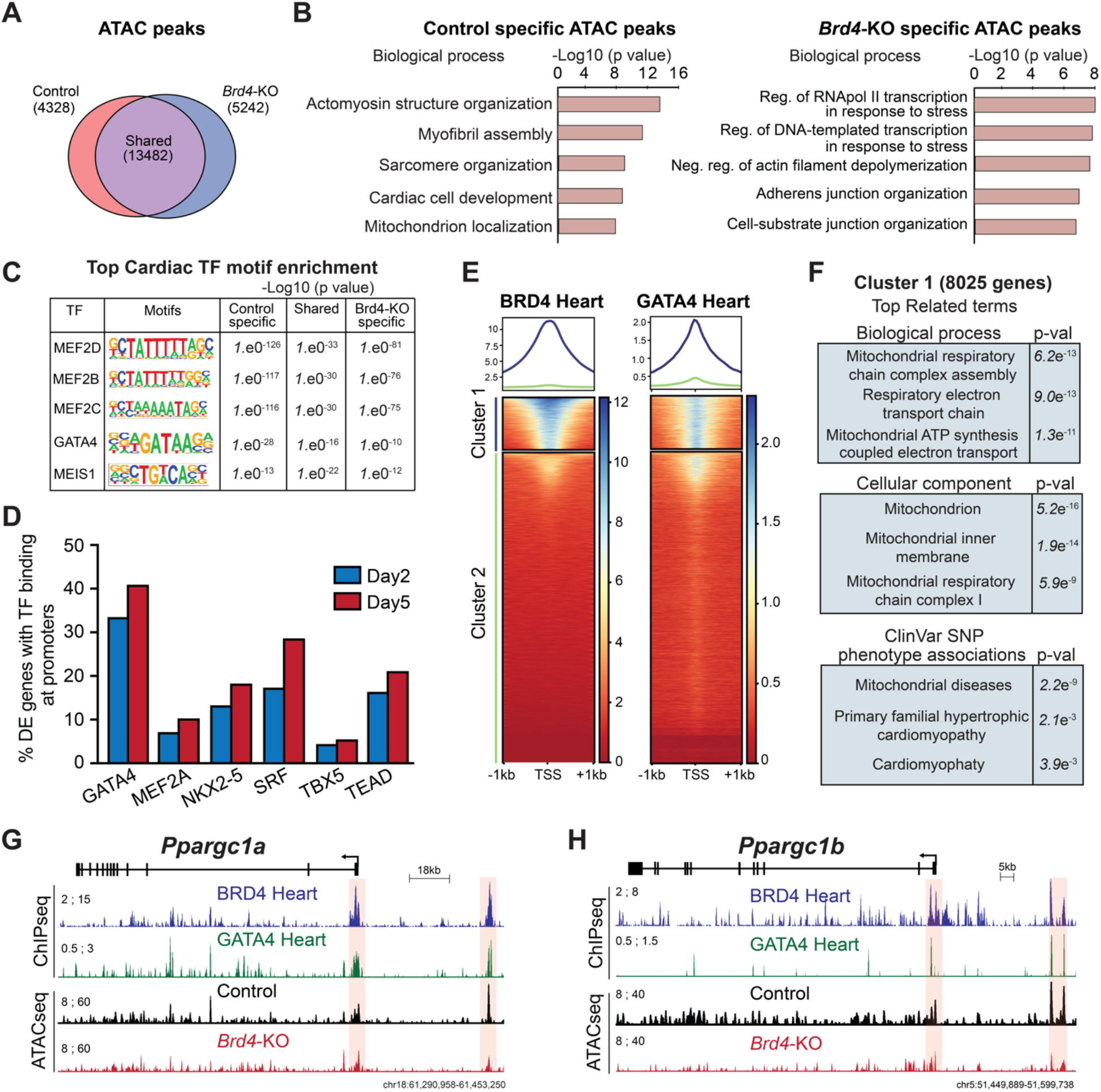
BRD4 and GATA4 co-occupy and regulate genes controlling mitochondrial homeostasis. **(A)** Venn diagram showing number of unique and shared accessible chromatin regions between Control and *Brd4*-KO samples. **(B)** Top selected categories identified from gene ontology analysis from Control and *Brd4*-KO specific ATAC regions. **(C)** Motif enrichment analysis for cardiac TFs in unique and shared accessible chromatin regions between Control and *Brd4*-KO samples. **(D)** Number of differentially expressed genes between Control and *Brd4*-KO at day 2 and day 5 occupied by cardiac TFs^32,33^ at their promoters (±1 TSS). **(E)** Heatmaps showing enrichment of BRD4 and GATA4 ChIP signal at gene promoters (±1 TSS, 55,386 mm10 annotated transcripts) ordered by BRD4 intensity identifies a cluster of strongly bound genes (Cluster 1, 8080 genes). **(F)** Gene ontology analysis of Cluster 1 genes identified enriched terms for biological processes, cellular components, and SNP-phenotype associations. **(G-H)** Track view of *Ppargc1a* and *Ppargc1b* genes showing sequencing reads mapping from BRD4 and GATA4 ChIPseq as well as Control and *Brd4*-KO ATACseq at day 5 post-tamoxifen treatment.

We were intrigued by the relative specificity of the transcriptional programs that were altered following CM-deletion of *Brd4*, which occupies regulatory regions throughout the genome. As BRD4 has previously been demonstrated to interact with sequence-specific TFs in other contexts^26–30^, we hypothesized that tissue-restricted TFs may facilitate preferential enrichment of BRD4 at specific genomic loci and contribute to the sensitivity of certain gene programs to BRD4 depletion. To test this hypothesis, we performed motif analyses^31^ of all the elements identified in our ATAC-seq and found enrichment for several cardiac TF motifs, including those for members of the MEF2 family, GATA4, and MEIS1 (**Figure 3C**). Interestingly, GATA4 was the only cardiac TF motif whose enrichment demonstrated a graded decrease in significance from those peaks unique to control CMs (lost in *Brd4*-KO CMs), shared between both conditions, and those unique to *Brd4*-KO CMs (gained in *Brd4*-KO CMs), suggesting that GATA4 may have preferential function in regions of active chromatin that were most sensitive to the presence of BRD4. Importantly, publicly available datasets defining the occupancy of cardiac TFs in adult heart tissue^32,33^ demonstrate that promoters of differentially expressed genes in *Brd4*-KO CMs are enriched in GATA4 occupancy as compared to other cardiac TFs (**Figure 3D**).

Given this enrichment for GATA4 binding sites in the differentially expressed genes of *Brd4*-KO CMs, we posited that BRD4 and GATA4 may co-occupy the promoters of these genes. To explore this hypothesis, we analyzed our previously performed BRD4 ChIP-seq^9^ and publicly available GATA4 ChIP-seq datasets obtained from adult myocardium^32^. We clustered the enrichment of BRD4 and GATA4 occupancy in regions proximal to the gene transcriptional start site (TSS, ±1 kb) of all annotated genes and found that genes separated into two clusters, one with strong occupancy of BRD4 and GATA4 (cluster 1; 8,080 genes) and a second with little or absent co-occupancy (cluster 2; 47,305 genes) (**Figure 3E**). Like our RNA-seq data, GO analysis of cluster 1 revealed genes for mitochondrial bioenergetics. Similar ontology analyses identified the mitochondrion as the most enriched cellular component, and genome-wide association studies linked single nucleotide polymorphisms (SNPs) from these genes with mitochondrial disease and cardiomyopathy (**Figure 3F**). These data indicate that BRD4 and GATA4 co-occupy promoter regions of genes involved in mitochondrial energy production in adult CMs.

Given the downregulation of *Ppargc1a* and *Ppargc1b* at day 2 after *Brd4* deletion, we specifically examined these loci for evidence of direct regulation by BRD4 and GATA4. Both of these genes exhibited strong enrichment for BRD4 and GATA4 at their promoters and were characterized by a putative upstream regulatory element that was also co-occupied by BRD4 and GATA4 (**Figure 3G,H**). Notably, both of these upstream DNA elements showed decreased accessibility in *Brd4*-KO CMs when compared with controls, suggesting that accessibility of these regulatory regions are BRD4-dependent (**Figure 3G,H**). Although HF can be associated with a secondary downregulation of CM metabolic genes, our findings suggest a model in which a BRD4-GATA4 module controls cardiac metabolic gene programs through a feed-forward mechanism, regulating both nodal upstream transcriptional coactivators of mitochondrial metabolism as well as downstream gene targets.

### BRD4 forms a complex with GATA4 in an BD-independent fashion

Given the striking colocalization of BRD4 and GATA4 at regulatory regions of BRD4-sensitive genes, we hypothesized that BRD4 and GATA4 may functionally co-regulate these targets. We began by testing the ability of BRD4 and GATA4 to transactivate an *Nppa* promoter-luciferase reporter construct in transient transfection assays. Transfection of GATA4 or BRD4 alone resulted in an approximate 3-fold or 3.5-fold activation of this reporter, respectively, while co-transfection of both GATA4 and BRD4 increased reporter activity by approximately 5.5-fold, consistent with functional co-regulation (**Figure 4A**).

**Figure 4:**
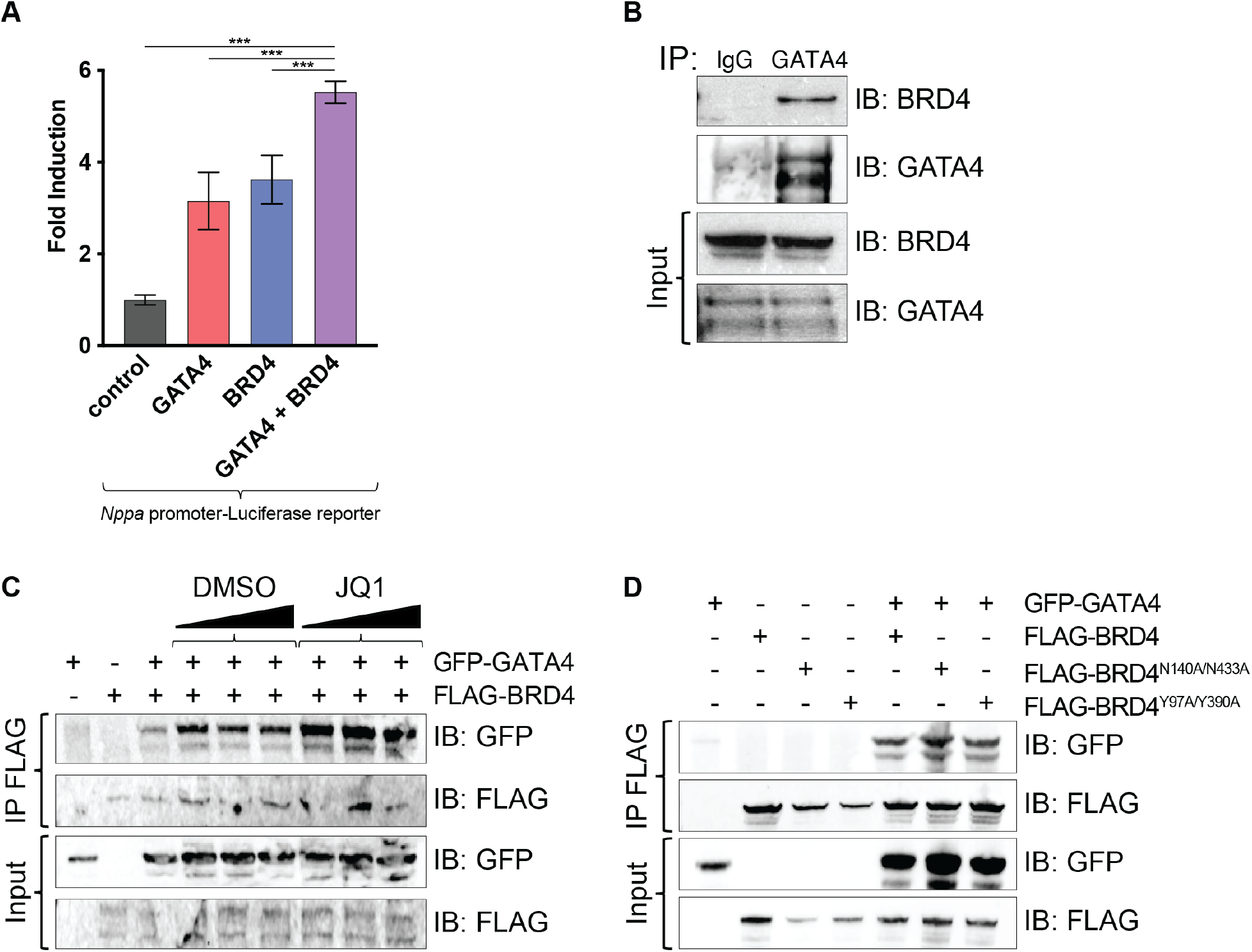
BRD4 physically interacts with GATA4 in a bromodomain-independent manner. **(A)** Gene reporter assay showing activation of luciferase reporter upon addition of plasmids encoding indicated proteins. Mean ± 1 SD shown. *** indicates p<0.00001. **(B)** Immunoprecipitation (IP) of endogenous protein from human cardiac progenitor cells using anti-GATA4 or anti-IgG antibody and immunoblotting (IB) with anti-BRD4 or anti-GATA4 antibody demonstrates endogenous BRD4 co-immunoprecipitates with GATA4 but not IgG. **(C)** Immunoprecipitation of FLAG-BRD4 overexpressed in HEK293 cells followed by immunoblotting with anti-GFP or anti-FLAG antibodies demonstrates GFP-GATA4 still co-immunoprecipitates with BRD4 even in the presence of increasing doses of JQ1 or DMSO as control. **(D)** Immunoprecipitation of FLAG-BRD4, FLAG-BRD4^N140A/N433A^, and FLAG-BRD4^Y97A/Y390A^ in HEK293 cells followed by immunoblotting with anti-GFP or anti-FLAG antibodies indicates co-immunoprecipitation of BRD4 mutants with GFP-GATA4.

Immunoprecipitation of endogenous GATA4 protein from human induced pluripotent stem cell-derived cardiac progenitor cell lysates (which express high levels of GATA4), followed by immunoblotting for BRD4, indicates that endogenous human GATA4 and BRD4 interact in CM progenitors (**Figure 4B**). When co-transfected in 293T cells, GFP-labelled GATA4 and FLAG-tagged BRD4 could also co-immunoprecipitate (**Figure 4C**). GATA4 contains four lysine residues in close proximity to its second zinc finger that are targets of p300-mediated acetylation^36^. These residues are conserved in GATA1, which has previously been demonstrated to bind to BRD3 in an acetyl-lysine dependent fashion via BRD3’s N-terminal BDs^37,38^. We explored if the BRD4-GATA4 complex was also dependent on the BDs of BRD4 by testing whether the complex could be disrupted by JQ1. In transfected 293T cells, increasing concentrations of JQ1 had no appreciable effect on the ability of BRD4 and GATA4 to interact (**Figure 4C**). The tertiary structure of the BRD4 BD bound to acetyl-lysine^11^ and JQ1^19^ has been extensively characterized with complementary biochemical and biophysical methods, revealing 2 amino acids in each BD that are particularly critical for mediating interactions with lysine-acetylated protein targets. The acetyl-lysine side chain is recognized by a central hydrophobic cavity that is anchored by a hydrogen bond with a conserved asparagine residue (N140 of BD1; N433 of BD2) while a second interaction is formed between the acetyl carbonyl oxygen atom and a conserved tyrosine residue (Y97 of BD1; Y390 of BD2) via a water molecule^39–42^. BRD4 mutant constructs harboring N140A and N433A mutations or Y97A and Y390A retained the ability to interact with GATA4 (**Figure 4D**), consistent with this interaction occurring independent of the BRD4 BDs. This observation is consistent with the disparity in cardiac functional consequences of CM-specific *Brd4* deletion and JQ1 treatment in mice.

## Discussion

In this report, we leveraged a newly developed mouse line harboring a conditional *Brd4* allele and unbiased transcriptomic and epigenomic assays to dissect the tissue-specific role of this transcriptional coactivator in adult CM homeostasis *in vivo*. Acute depletion of BRD4 in adult CMs caused rapid onset of systolic HF within 5 days, characterized by a dramatic decrease in LV systolic function and LV chamber dilation, leading to 100% lethality by 20 days. Transcriptomic profiling of CMs from *Brd4*-KO mice at day 5 revealed robust and relatively specific downregulation of gene programs important for mitochondrial energetics and homeostasis. Integrated analysis of BRD4 dependent gene expression and chromatin accessibility in adult CMs identified that GATA4 binding sites were preferentially enriched in the regulatory regions of BRD4-dependent genes. Our results reveal that BRD4 and GATA4 co-localize across the genome at loci relevant to mitochondrial bioenergy production and identify a novel BD-independent protein complex formation between BRD4 and this key cardiac TF. These findings highlight an unexpected role for both BRD4 and GATA4 in regulating a metabolic gene expression program in adult CMs.

BET proteins have emerged as intensely pursued pharmacologic targets in cancer and several chronic diseases, including HF^7,8,14,43–45^. In animal models of HF, small molecule BET BD inhibitor therapy improves cardiac function, decreases fibrosis, and attenuates transactivation of inflammatory and pro-fibrotic gene programs^9,10,20^. Clinical grade BET BD inhibitors are currently in trials for the treatment of a variety of malignancies^14^. Although chemically diverse, these compounds share a common structural motif that reversibly binds the N-terminal BDs of all BET family members, thereby transiently displacing them from their endogenous substrates. BRD2, BRD3, and BRD4 are all ubiquitously expressed and the relevant tissue compartments in which they function *in vivo* remains unknown. Small molecule proteolysis targeting chimeras (PROTACs) that degrade BET proteins represent a closer equivalent to genetic deletion^46–48^, but these again are neither cell specific nor selective for individual BET isoforms. Therefore, none of these current pharmacologic approaches offer any insight into the cell-type and isoform specificities associated with beneficial responses of these compounds in chronic disease models. Our data show that acute BRD4 depletion in CMs of the adult mouse leads to rapid-onset systolic HF and mortality, highlighting a sharp contrast to the salubrious effects of small molecule BET BD inhibitors in treating adult mice with HF. These data are consistent with earlier transcriptomic characterization of JQ1 treated hearts and suggest that the dominant cellular targets of these compounds in the context of HF are non-CMs, such as fibroblasts and immune cells^20,49^. Further dissection of these cell compartment- and gene-specific effects in the context of HF using conditional mice for each BET allele will be a fruitful area of future study. While small molecule BET BD inhibitors and conditional gene deletion approaches manipulate BET proteins by very different mechanisms, our findings underscore the importance of understanding the specific cell-types that mediate the beneficial effects and potential liabilities of BET BD inhibitor therapy.

Our studies also suggest a previously unrecognized role for GATA4, perhaps one of the most widely studied TFs in cardiac biology, in CM metabolism and mitochondrial homeostasis. GATA4 is a well-established lineage determining TF for CMs whose role in cardiac development and human congenital heart disease has been extensively studied by our group and others^50,51^. As ectopic *Gata4* expression (along with *Mef2c* and *Tbx5*; GMT) can induce conversion of cardiac fibroblasts into induced CMs^52–54^, much effort has also focused on the molecular mechanisms by which GATA4 regulates cellular identity^55,56^. In the adult heart, GATA4 plays an important role in stress responsiveness and regulation of pro-angiogenic genes^57,58^. Our findings suggest that the preference of BRD4 to occupy certain loci may be mediated, in part, through an interaction with GATA4. Functionally, our data indicate that BRD4 governs mitochondrial homeostasis in the adult heart. Loss of *Brd4* in CMs results in clear loss of mitochondrial structure and organization. Our data indicate that a BRD4-GATA4 co-regulatory module governs mitochondrial gene expression programs by a tiered, feed-forward mechanism regulating both nodal upstream regulators of these transcriptional outputs along with their downstream effectors.

Cardiac contractile function is highly dependent on the ability of working CMs to sustain very high rates of mitochondrial oxidative metabolism and efficiently transfer energy^59,60^. Human cardiomyopathies caused by loss-of-function variants in key mitochondrial genes underscore the direct link between mitochondrial dysfunction and contractile failure^61,62^. Consistent with these findings, deletion of key transcriptional regulators of mitochondrial homeostasis in murine CMs, including PGC1-α and ERRα, cause cardiomyopathy and HF^34,35,63^. Electron microscopy of *Brd4*-KO hearts also revealed myofiber degeneration and sarcomere disarray. This phenotype may be a downstream consequence of mitochondrial dysfunction, which can lead to pleiotropic defects in cellular homeostatic processes, including protein quality control, redox balance, and calcium handling. In parallel, BRD4 may also play a primary role in coactivating key genes that are critical for sarcomere assembly and maintenance. In the developing heart, GATA4 is required for cardiac morphogenesis and directly activates sarcomere genes during cardiac lineage commitment^64^, raising the possibility that the BRD4-GATA4 axis in adult CMs may regulate sarcomere genes in addition to mitochondrial programs. Dissecting the mechanisms by which BRD4 deficiency leads to both mitochondrial dysfunction and sarcomere disarray in CMs will be an important area of future investigation.

Our experiments have also revealed critical new knowledge about a commonly used reagent, the *Myh6-MCM* transgenic mouse^23^. This mouse strain has become the *de facto* standard tool for inducible CM-targeted deletion of conditional alleles in adult CMs. This transgene expresses a Cre recombinase flanked by mutated estrogen receptor ligand-binding domains insensitive to endogenous estrogen (but responsive to exogenously administered tamoxifen) under the influence of the ɑ-myosin heavy chain (*Myh6*) promoter. Prior reports have demonstrated a transient myopathy associated with Cre nuclear translocation in these animals following tamoxifen administration and warned of potential confounding in assessment of acute phenotypes that result from ablation of any gene of interest^24^. Here, we define the first transcriptomic characterization of this transient myopathy (**Figure S4**) at days 2 and 5 after tamoxifen administration. Hence, accounting for gene expression changes upon induction of this transgene will be critical in the interpretation of results generated using these mice, providing a major resource for the field.

Apart from the potential clinical application of BET BD inhibitors for therapeutic purposes, compounds like JQ1 have emerged as powerful tools for investigating enhancer biology^65^. Indeed, much of the field focuses on the BD-mediated functions of these proteins. In addition to acetylated histones, BET proteins have been found to bind acetylated TFs and modulate their transcriptional output. For example, a BD-dependent interaction between BRD3 and GATA1 is important for erythroid maturation^37^. Likewise, BD-dependent interactions between BRD4 and p65/RelA-K310ac have been implicated in innate immunity^26^ and similar interactions with several hematopoietic TFs (including PU.1, FLI1, and ERG) are essential for acute myeloid leukemia pathogenesis^30^. The advent of small molecules with a selectivity for BD1 vs. BD2 have stimulated further interest in dissecting the relative contribution of each of these domains in a variety of cellular contexts^66,67^. However, the BETs are large proteins that can scaffold complexes via multiple domains outside of their N-terminal BDs. For example, the BRD4 C-terminal domain interacts with the PTEFb complex^16,17^ and the BRD4 extra-terminal domain interacts with the histone methyltransferase NSD3^68,69^. Unbiased protein-protein interaction screens for BET family members have been performed both in the presence and absence of JQ1, demonstrating a number of BD-independent interacting partners for each of the ubiquitously expressed BET family members^70^. Our data identify a novel, BD-independent interaction between BRD4 and GATA4, suggesting a mechanism by which a broadly expressed chromatin coactivator can be preferentially targeted to specific genomic loci by associating with a tissue-enriched DNA-binding TF. JQ1-mediated inhibition *in vivo* would not be expected to disrupt the BRD4-GATA4 interaction, consistent with the differences seen between the therapeutic effects of JQ1 and the deleterious consequences of *Brd4* deletion *in vivo*. This work in CMs highlights a critical need to better understand structure-function relationships of BRD4 outside of the BDs, findings which will provide general insight into the molecular underpinnings of cell-specific gene regulation and inform novel therapeutic approaches for HF and other cardiovascular diseases.

## Acknowledgments

We thank the Srivastava and Jain laboratories for critical discussions and feedback; the Penn CVI Histology group for help with histology; the Penn Electron Microscopy Core for assistance with sectioning and imaging of electron micrographs; and the Gladstone Genomics Core for preparing and sequencing the RNA-seq libraries.

## Sources of Funding

A.P. is supported by the Tobacco-Related Disease Research Program (578649), A.P. Giannini Foundation (P0527061), and Michael Antonov Charitable Foundation Inc. M.A. is supported by the Swiss National Science Foundation (P400PM_186704). R.L.S. is supported by NIH F31 HL147416. S.M.H. was supported by NIH R01 HL127240. R.J. is supported by the Burroughs Wellcome Career Award for Medical Scientists, Gilead Research Scholars Award, NSF CMMI-1548571, and NIH R01 HL139783. D.S. is supported by NIH P01 HL098707, NIH R01 HL057181, and by the Roddenberry Foundation, the L.K. Whittier Foundation, and the Younger Family Fund. This work was also supported by NIH/NCRR grant C06 RR018928 to the Gladstone Institutes.

## Disclosures

D.S. is scientific co-founder, shareholder and director of Tenaya Therapeutics. S.M.H. is an executive, officer, and shareholder of Amgen, Inc. and is a scientific co-founder and shareholder of Tenaya Therapeutics.

---

**Supplementary Figure 1:**
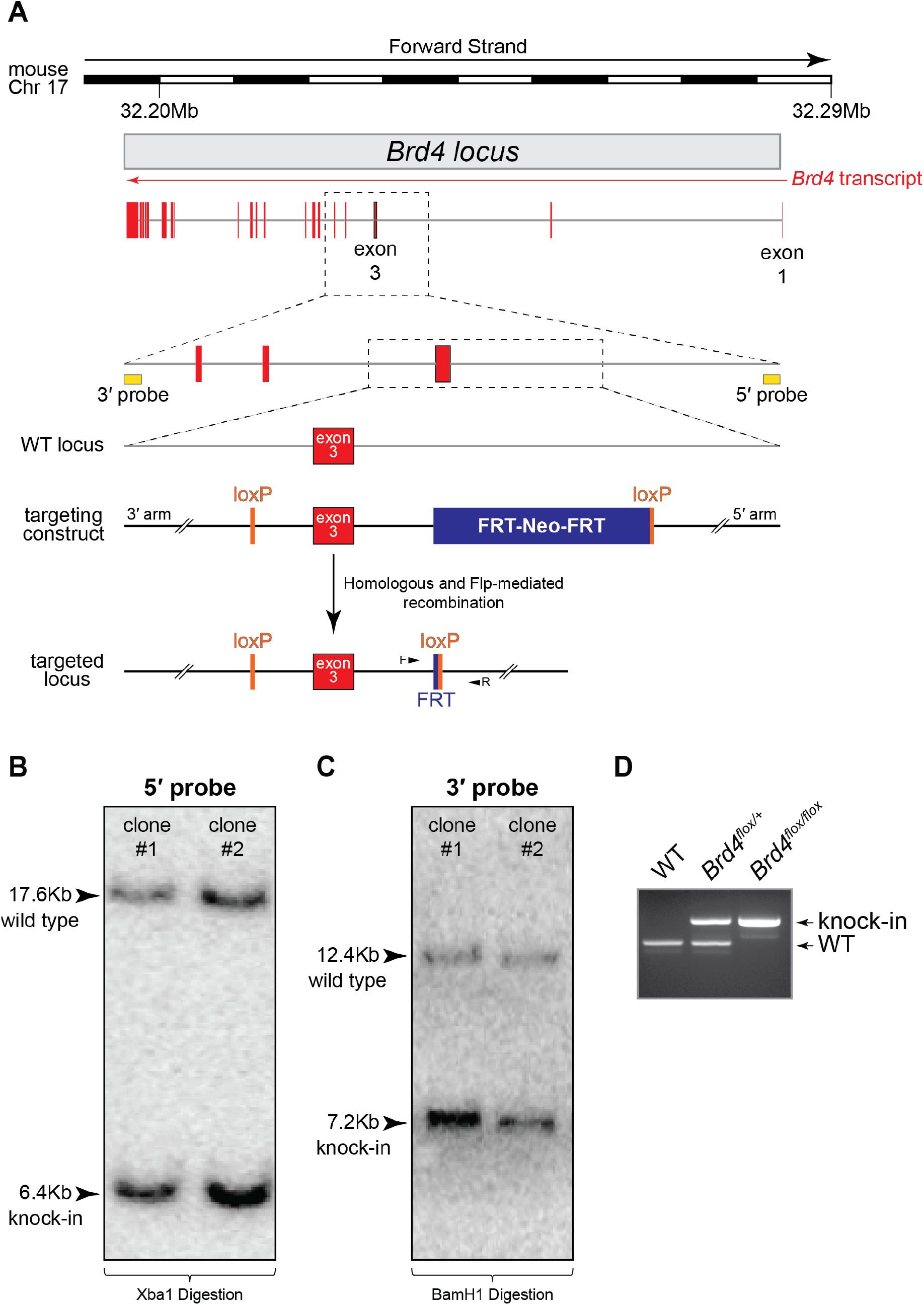
Generation of a *Brd4* conditional allele (*Brd4^flox^*). **(A)** Targeting strategy to flank *Brd4* exon 3 (containing the canonical translation start site) with loxP sites. **(B-C)** Southern blotting using both 5′ (B; wild type band 17.6Kb, knock-in band 6.4Kb) and 3′ (C; wild type band 12.4Kb, knock-in band 7.2Kb) probes confirms appropriate targeting of mouse embryonic stem cells (images from two representative clones displayed). **(D)** Representative image of PCR genotyping assay used for *Brd4^flox^* allele that resolves an 82 bp difference (representing the residual FRT scar and 5′ loxP site) between the wild type and knock-in allele.

**Supplementary Figure 2:**
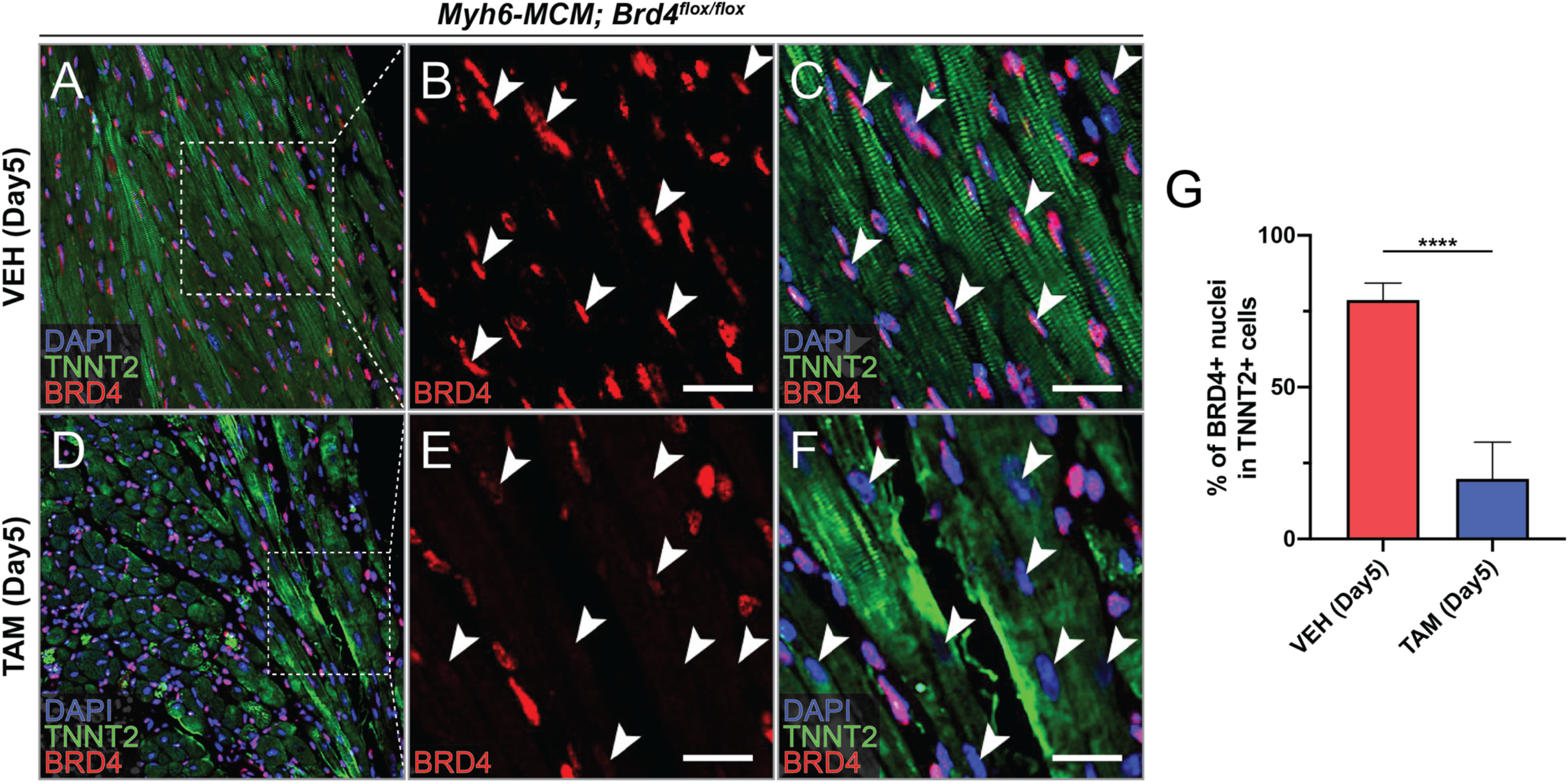
Immunohistochemical analysis of BRD4 in *Brd4*-KO and control hearts. **(A-F)** Sections stained with DAPI (blue), TNNT2 (green), and BRD4 (red) in vehicle (VEH; A-C) or tamoxifen treated (TAM; D-F) *Myh6-MCM; Brd4^flox/flox^* mice at 10-weeks of age. **(G)** Quantification of percentage of TNNT2+ cells with BRD4+ nuclei. Mean ± 1 SD shown, n=4 sections quantified. *** indicates p<0.0001. Scale bars = 20 μm.

**Supplementary Figure 3:**
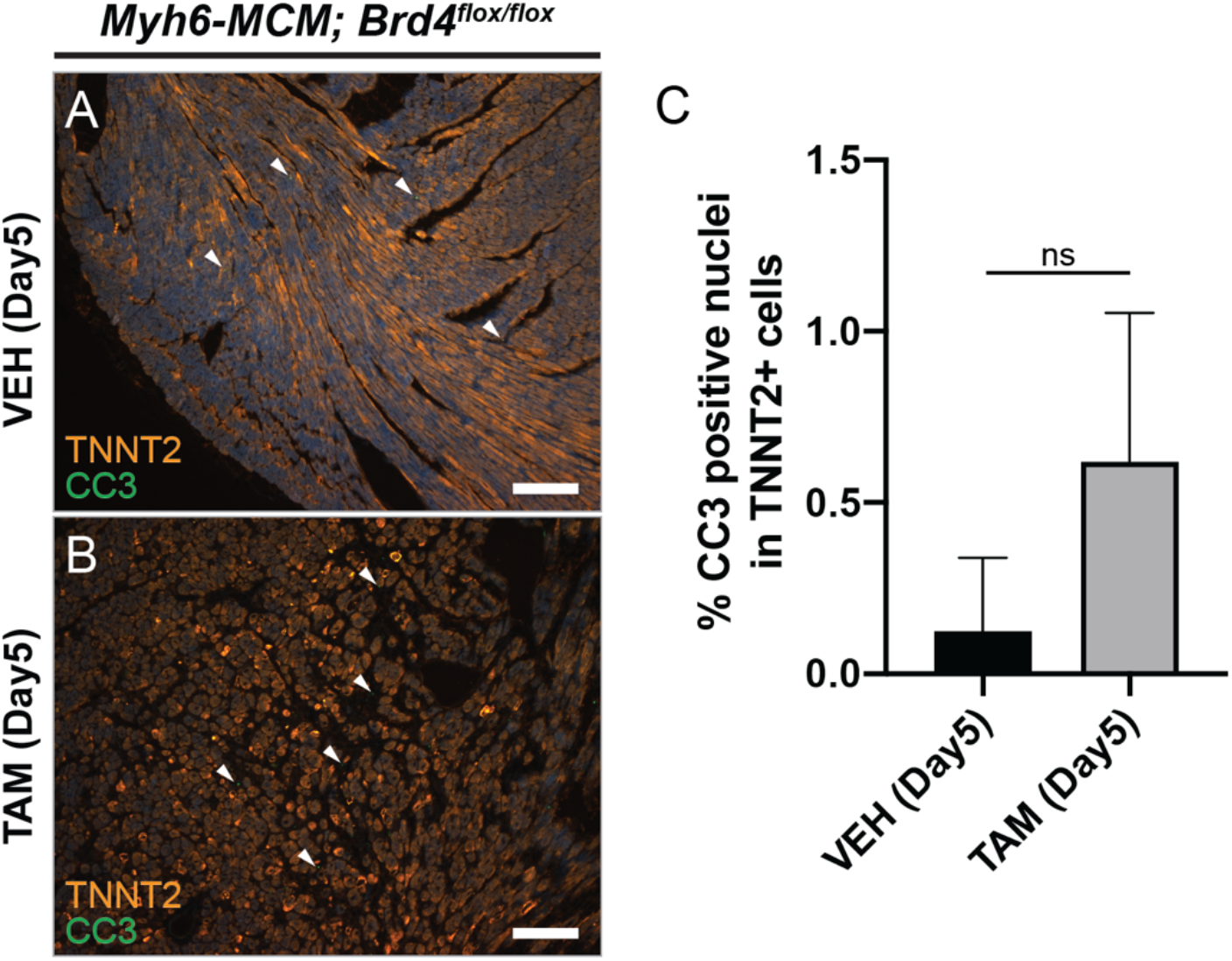
Cleaved caspase 3 staining in *Brd4*-KO and control hearts. **(A,B)** Sections from 10-week old *Myh6-MCM; Brd4^flox/flox^* mice treated with vehicle (VEH; A) or tamoxifen (TAM; B) stained with cleaved caspase 3 (CC3; green) and TNNT2 (orange). Mean ± 1 SD shown, n=3 sections quantified. Scale bars = 100 μm.

**Supplementary Figure 4:**
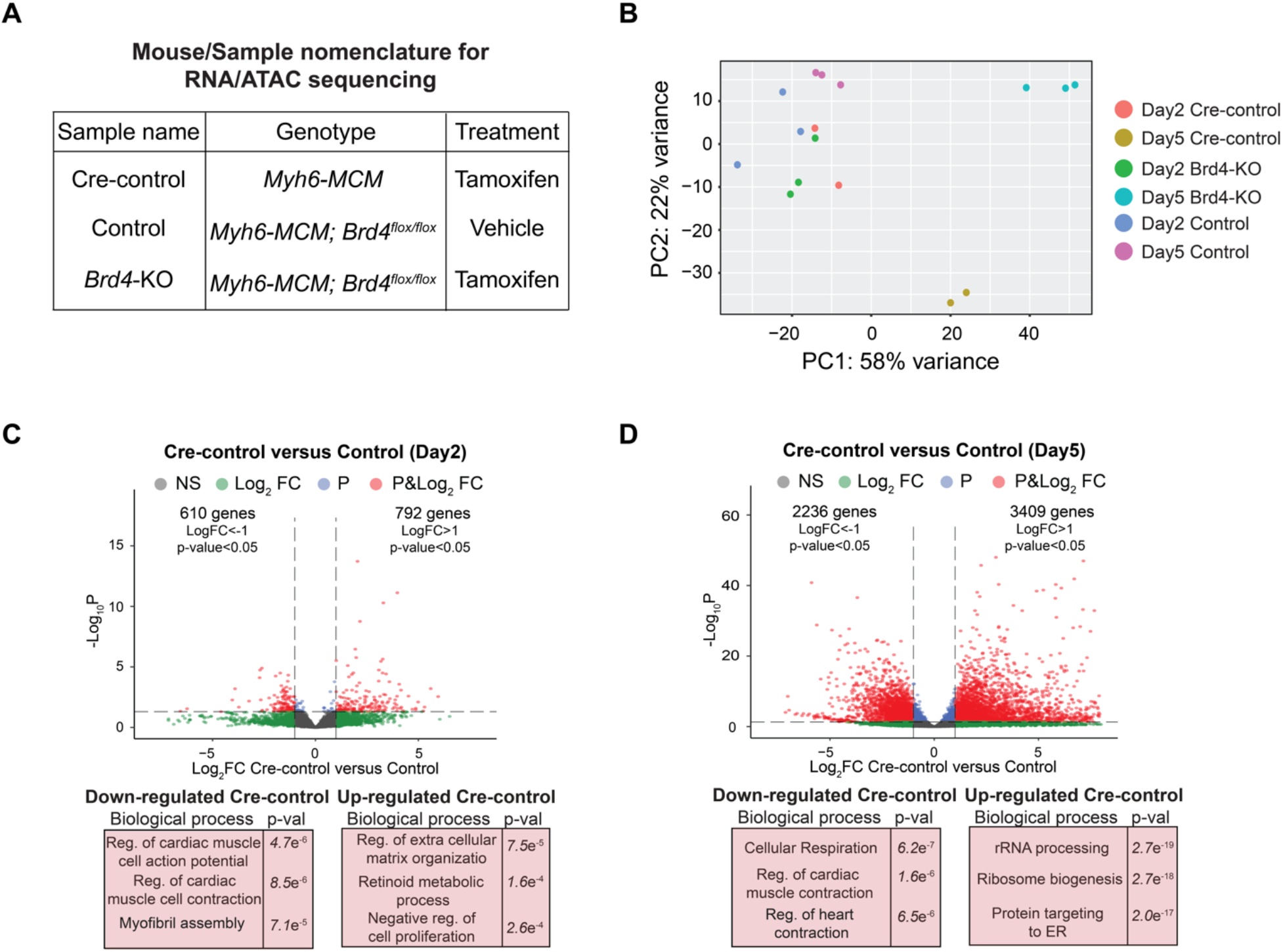
Gene expression changes following tamoxifen-mediated Cre nuclear localization in Cre-control adult cardiac myocytes. **(A)** Table indicating naming convention and genotypes of various samples used in this study. **(B)** PCA plot of individual replicates of each genotype used in RNA-seq analyses. **(C-D)** Volcano plots of differential gene expression in Cre-control vs. Control at day 2 (C) and day 5 (D) with associated enriched terms from gene ontology analysis.

**Supplementary Figure 5:**
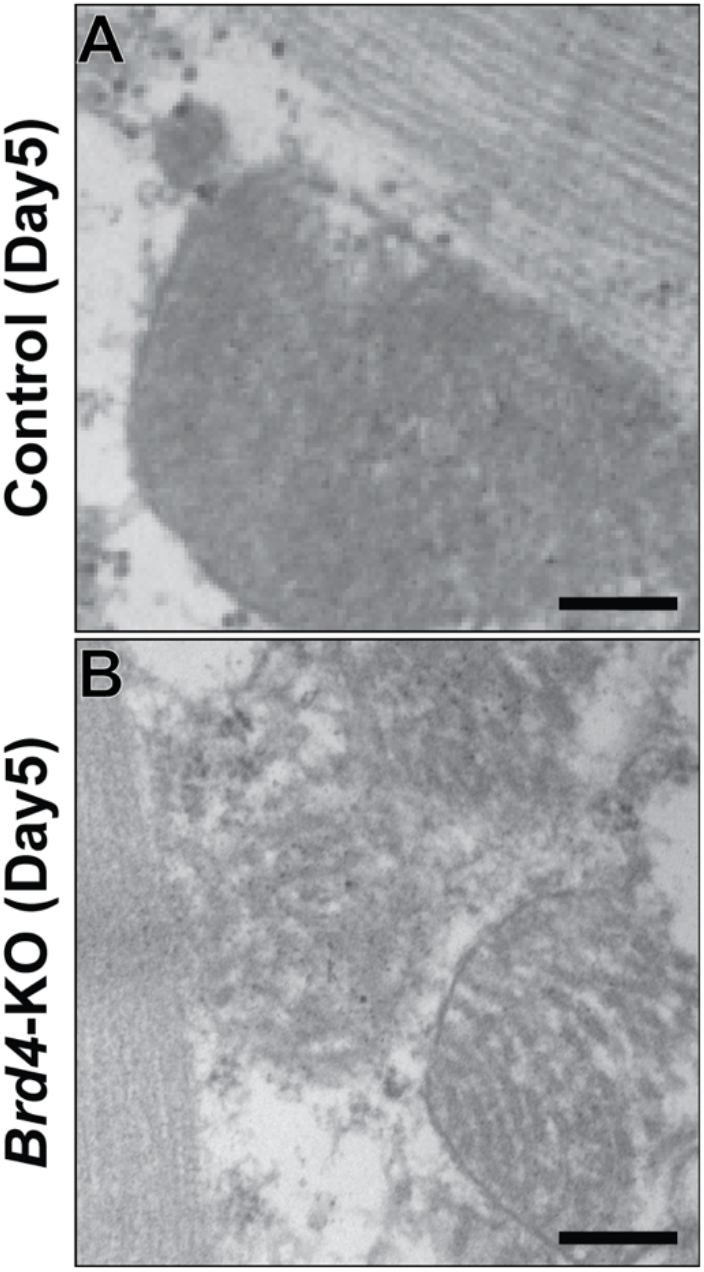
Electron micrographs of mitochondria in *Brd4*-KO and control hearts. **(A, B)** Electron micrographs of cardiac tissue from 10-week old *Myh6-MCM; Brd4^flox/flox^* mice treated with vehicle (VEH; A) or tamoxifen (TAM; B) highlighting the loss of normal mitochondrial morphology. Scale bars = 100 nm.

**Supplementary Tables 1-4:**

RNA-seq data at Day 2 or Day 5 in *Myh6-MCM; Brd4^flox/flox^* animals treated with tamoxifen (*Brd4*-KO) or vehicle (control) and RNA-seq data at Day 2 or Day 5 of Cre control (*Myh6-MCM*) versus control.

## Methods

### RNA sequencing, mapping, and quantification

Paired-end Poly(A)-enriched RNA libraries were prepared with the ovation RNA-seq Universal kit (NuGEN; strand specific). High-throughput sequencing was done using a PE75 run on a NextSeq 500 instrument (Illumina). Reads were mapped to the mm10 reference mouse genome using STAR (v 2.7.3a) and assigned to Ensembl genes. After read quality control, we obtained quantifications for 38,293 genes in all 16 samples (2 Day2 Cre-control, 2 Day5 Cre-control, 3 Day2 Control, 3 Day5 Control, 3 Day2 *Brd4*-KO and 3 Day5 *Brd4*-KO).

### Differential gene expression and pathway enrichment analysis

In order to identify genes differentially expressed in Cre-control, Control, and *Brd4*-KO samples we quantified gene expression using raw counts and performed differential expression gene testing with DESeq2^71^ (v.1.24.0 R package) using default settings. Statistical significance was set at 5% false discovery rate (FDR; Benjamini-Hochberg). Functional enrichment gene-set analysis for GO (Gene Ontology) terms was performed using Enrichr^72^. Heat maps were generated using the Bioconductor package pheatmap (v.1.0.12) using rlog-transformed counts (values shown are rlog-transformed and row-normalized). Volcano plots were generated using the Bioconductor package EnhancedVolcano (v.1.2.0).

### Comparison of RNA-seq datasets

To compare our RNA-seq results to the transcriptional changes associated with JQ1-mediated BET BD inhibition, we used published expression profiles of JQ1- and vehicle-treated sham-operated mouse hearts (GSE96561). The raw counts were analyzed using DEseq2 (v1.22.2 R package), and the correlation of the results were done by representing the Log2FC values from each study using ggplot2 (v3.2.0 R package). Data was taken from three biological replicates in each condition.

### Electron microscopy

Tissues for electron microscopic examination were fixed with 2.5% glutaraldehyde and 2.0% paraformaldehyde in 0.1M sodium cacodylate buffer (pH7.4) overnight at 4°C. After subsequent buffer washes, the samples were post-fixed in 2.0% osmium tetroxide with 1.5% K3Fe(CN)6 for 1 hour at room temperature, and rinsed in distilled water. After dehydration through a graded ethanol series, the tissues were infiltrated and embedded in EMbed-812 (Electron Microscopy Sciences, Fort Washington, PA). Thin sections were stained with uranyl acetate and SATO lead and examined with a JEOL 1010 electron microscope fitted with a Hamamatsu digital camera and AMT Advantage NanoSprint500 software.

### Mice

All mouse manipulations were performed in accordance with protocols approved by the Institutional Animal Care and Use Committee (IACUC) following guidelines described in the US National Institutes of Health Guide for the Care and Use of Laboratory Animals.

*Myh6-MCM* mice have been described previously^23^. *Brd4^flox^* mice wre produced by targeting C57BL/6 ES cells with a targeting vector designed to flank exon 3 (containing the canonical *Brd4* ATG) with loxP sites. The selection strategy included a FRT flanked neomycin resistance cassette that, after excision, leaves a single FRT site within intron 2. *Brd4^flox^* mice were genotyped using the polymerase chain reaction (PCR) primers listed below which produce a 311 bp wild-type band and a 414 bp knock-in band (Figure S1D). All mice were maintained on a C57BL/6 background.

Brd4flox_01F: 5′-GAAAGAGAAGAAGCTAACTGGC
Brd4flox_02R: 5′-GAGCAAGTATATTGGAGGGGAG.

### Mouse echocardiography

Echocardiography was performed blindly with the Vevo 770 High-Resolution Micro-Imaging System (VisualSonics) with a 15-MHz linear-array ultrasound transducer. The left ventricle was assessed in both parasternal long-axis and short-axis views at a frame rate of 120 Hz. End-systole or end-diastole were defined as the phases in which the left ventricle appeared the smallest and largest, respectively, and used for ejection-fraction measurements. To calculate the shortening fraction, left-ventricular end-systolic and end-diastolic diameters were measured from the left-ventricular M-mode tracing with a sweep speed of 50 mm/s at the papillary muscle. B-mode was used for two-dimensional measurements of end-systolic and end-diastolic dimensions. Imaging and calculations were done by an individual who was blinded to the treatment applied to each animal and code was broken only after all data acquired.

### Histology

Isolated hearts were fixed in 2% paraformaldehyde (4°C overnight), dehydrated through an ethanol series, embedded in paraffin, and sectioned. Antibodies used for immunohistochemistry were: Brd4 (rabbit, Bethyl 00396) and Cleaved caspase 3 (rabbit, Cell Signaling 9664). Hematoxylin and Eosin staining was performed using standard protocols. Sections were imaged on a Nikon Eclipse 80i fluorescence microscope or Leica DMi8 inverted fluorescence microscope.

### ATAC-sequencing, library preparation, and analysis

CM samples were prepared for ATAC-seq as previously described^25^. Aliquots of 50,000 cells were were lysed with 3 mL of chilled lysis buffer (3.75mM PIPES, 450mM KCl, 1% NP-40, 1% Tween-20, 1% Triton X-100 in water; pH 7.3) for 10 minutes. Nuclear pellets were transposed with 25 μL of Tagment DNA Buffer, 2.5 μL of Tagment DNA Enzyme (Nextera Sample Prep Kit from Illumina; #FC-121-1030), and 22.5 μL of nuclease-free water. Samples were incubated at 37°C for 60 min and stored at 20°C. Transposed samples were purified using the QIAGEN MinElute Reaction Cleanup Kit (28204) and amplified using 25 μL of NEBNext High Fidelity 2x PCR Master Mix, 1.25 mM Nextera custom primers with unique barcodes, and nuclease-free water. Samples were amplified using the following PCR conditions: 72°C for 5 min; 98°C for 30 s; and cycled at 98°C for 10 s, 63°C for 30 s and 72°C for 1 min. Half of each sample was amplified for 12 cycles, purified, and assessed by Bioanalyzer (Agilent) for library quality. Sample concentration was quantified by Qubit (Invitrogen) prior to pooling. Pooled samples were sequenced using a PE75 run on a NextSeq 500 instrument (Illumina). Alignment to the mm10 reference genome was performed using Bowtie 2.2.4. Peaks were called using macs2 callpeak with options: “-p 0.1–nomodel–shift 100–extsize 200-B–SPMR–call-summits”. Peaks concordant between the two replicates were considered for further analysis. Motif enrichment analysis was performed using HOMER^31^.

### Analysis of publicly available ChIP-seq data

Occupancy analysis for BRD4 and GATA4 was done using publically available ChIP-seq datasets (GSE52123 and GSE124008, respectively). Deeptools v.3.3.1 was used to generate the aggregation and heatmap plots centered at the TSS of mm10 gene annotations (GRCm38.p6). Analysis of transcription factor representation in our list of DEGs from Day2 and Day5 was done using published data (GSE124008) after filtering for peaks within 5Kb of the closest annotated gene.

### Immunoblotting

Adult CMs were isolated via Langendorff perfusion of *Myh6-MCM; Brd4^flox/flox^* treated with TAM (75 μg/g/day) or VEH (corn oil) for 5-days. Whole cell extracts were prepared by lysis in RIPA buffer supplemented with protease and phosphatase inhibitors (Roche). Protein concentration was quantified by BCA assay (Thermo Fisher; 23225). Lysates were diluted in 4X LDS sample buffer (Invitrogen), boiled at 95°C for 5 min, resolved on a 3-8% Tris-Acetate SDS PAGE gel (Invitrogen), and transferred onto a PVDF membrane. Membranes were blocked with 5% milk in TBST for 1 hour at room temperature and incubated in primary antibody at the indicated dilution overnight at 4°C. Appropriate secondary HRP-conjugated antibody was added for 1 hour at a dilution of 1:5000 followed by detection with ECL Prime Western Blotting Detection Reagent (GE Life Sciences; RPN2232) and exposure to autoradiography film at various time intervals or by digital imaging (LI-COR Odyssey). Antibodies used in this study: rabbit anti-Brd4 (Abcam ab128874; 1:1000) and mouse anti-vinculin (Sigma-Aldrich V9131; 1:1000).

### Luciferase reporter assay

GATA4–BRD4 transcriptional synergy reporter assay was performed using the pANF638L vector^73^. Briefly, HeLa cells were cultured in 24-well plates at 10^5 cells per well and transfected within 24 hrs of seeding. Cells were co-transfected with 200 ng of pANF638L vector and 20 ng of pRL-TK (a Renilla luciferase control vector; Promega) in 2.4 μL FuGENE HD (Promega) and 43 μL Opti-MEM (Thermofisher). The transfection mix was aliquoted in 4 tubes (1 per condition) and the following conditions were prepared: 1.) Control: 600ng of empty vector (EV); 2.) GATA4: 200ng of GFP-GATA4 vector plus 400ng EV; 3.) BRD4: 400ng FLAG-BRD4 vector plus 200ng EV and 4.) GATA4+BRD4: 200ng of GFP-GATA4 vector with 400ng FLAG-BRD4 vector. Media was changed 24 hrs after transfection and cells collected 48 hrs following transfection. Samples were processed with the Dual Luciferase Assay System (E1960, Promega) following manufacturer’s instructions and measured with a luminometer (SpectraMax i3). The GATA4–BRD4 transcriptional synergy was tested in three technical replicates in 3 independent experiments.

### Co-Immunoprecipitation in HEK 293T cells

HEK293T cells were plated at 60% confluence in DMEM with 10% FBS 12–15 hrs before transfection in 100 mm plates. Cells were transfected with transfection mix containing 50 μL FuGENE HD (Promega) and 1000 μL Opti-MEM (Thermofisher) along with 2 g GFP-GATA4 vector, 8 g of FLAG-BRD4 vector, and (when only one plasmid indicated) empty vector to 10 g of DNA. Media was changed 24 hrs after transfection and cells collected at 48 hrs. Cell pellets from a confluent 100 mm plate were lysed with 1 ml of cell lysis buffer [20 mM Tris-HCl pH 8, 85 mM KCl, 0.5% NP-40, and protease inhibitors], incubated for 10 min at 4°C, and centrifuged at 2,500 rpm for 5 min at 4°C to pellet nuclei. Supernatants were removed and pellets resuspended in 300 μL of nuclei extraction buffer (NE buffer) [20 mM HEPES pH 7.4, 0.5 M NaCl, 2 mM MgCl2, 1 mM CaCl2, 0.5% NP-40, 110 mM K-Acetate, 1 μM ZnCl2, benzonase, and protease/phosphatase inhibitors] and incubated for 30 min at 4°C. Nuclear enriched lysates were centrifuged at maximum speed 10 min at 4°C and supernatants transferred to a new tube and diluted with 600 μL (1:3) of IP dilution buffer [20 mM HEPES pH 7.9, 1 mM EDTA, 0.02% NP-40, and protease/phosphatase inhibitors]. Prior to IP, 30 μL was saved as input. 25 μL of magnetic Protein G Dynabeads were coated with 2 μg of M2 anti-FLAG antibody (F1804, Sigma) for 1 hr at 4°C. For IP, 25 μL of FLAG-coated beads were added to 150 g of nuclear-enriched total protein, incubated overnight at 4 °C with agitation, and washed four times in IP dilution buffer. Beads were boiled for 10 min at 95 °C in 20 μl of sample buffer. Extracts and immunoprecipitates were examined by SDS–PAGE and blotted with antibodies for the indicated targets.

### Co-Immunoprecipitation in cardiac progenitors cells

Human induced-pluripotent stem cells were differentiated into cardiomyocytes using Wnt pathway modulation^74^ and 4 x 12-well plates were collected at cardiac progenitor stage (day 6 of differentiation), pooled together, and snap frozen in liquid nitrogen. Cardiac progenitor cell pellets were lysed with 1 ml of cell lysis buffer [20 mM Tris-HCl pH 8, 85 mM KCl, 0.5% NP-40, and protease inhibitors], incubated 10 min at 4°C, and centrifuged at 2,500 rpm for 5 min at 4°C to pellet nuclei. Supernatant was removed and pellets resuspended in 600 μL of nuclei extraction buffer (NE buffer) [20 mM HEPES pH 7.4; 0.5 M NaCl, 2 mM MgCl2, 1 mM CaCl2, 0.5% NP-40, 110 mM K-Acetate, 1 μM ZnCl2, benzonase, and protease/phosphatase inhibitors], and incubated for 30 min at 4°C. Nuclear enriched lysates were centrifuged at maximum speed for 10 min at 4°C and supernatants transferred to a new tube and diluted with 1200 μL (1:3) IP dilution buffer [20 mM HEPES pH 7.9, 1 mM EDTA, 0.02% NP-40, and protease/phosphatase inhibitors]. Prior to IP, 30 μL was saved as input. 50 μL of magnetic Protein G Dynabeads were coated with 4 g of anti-GATA4 (G4) antibody (sc-25310X, Santa Cruz) for 1 hr at 4°C. For IP, 50 μL of GATA4-coated beads were added to 2 mg of nuclear-enriched total protein, incubated overnight at 4 °C with agitation, and washed three times in IP dilution buffer. Beads were boiled for 10 min at 95 °C in 20 μl of sample buffer. Extracts and immunoprecipitates were examined by SDS–PAGE and blotted with antibodies for the indicated targets.

## References

1. Virani SS, Alonso A, Benjamin EJ, Bittencourt MS, Callaway CW, Carson AP, Chamberlain AM, Chang AR, Cheng S, Delling FN, Djousse L, Elkind MSV, Ferguson JF, Fornage M, Khan SS, Kissela BM, Knutson KL, Kwan TW, Lackland DT, Lewis TT, Lichtman JH, Longenecker CT, Loop MS, Lutsey PL, Martin SS, Matsushita K, Moran AE, Mussolino ME, Perak AM, Rosamond WD, Roth GA, Sampson UKA, Satou GM, Schroeder EB, Shah SH, Shay CM, Spartano NL, Stokes A, Tirschwell DL, VanWagner LB, Tsao CW, American Heart Association Council on Epidemiology and Prevention Statistics Committee and Stroke Statistics Subcommittee. Heart Disease and Stroke Statistics-2020 Update: A Report From the American Heart Association. Circulation. 2020;141:e139–e596.

2. Timmis A, Townsend N, Gale CP, Torbica A, Lettino M, Petersen SE, Mossialos EA, Maggioni AP, Kazakiewicz D, May HT, De Smedt D, Flather M, Zuhlke L, Beltrame JF, Huculeci R, Tavazzi L, Hindricks G, Bax J, Casadei B, Achenbach S, Wright L, Vardas P, European Society of Cardiology. European Society of Cardiology: Cardiovascular Disease Statistics 2019. Eur Heart J. 2020;41:12–85.

3. Burnett H, Earley A, Voors AA, Senni M, McMurray JJV, Deschaseaux C, Cope S. Thirty Years of Evidence on the Efficacy of Drug Treatments for Chronic Heart Failure With Reduced Ejection Fraction: A Network Meta-Analysis. Circ Heart Fail [Internet]. 2017;10. Available from: http://dx.doi.org/10.1161/CIRCHEARTFAILURE.116.003529

4. van Berlo JH, Maillet M, Molkentin JD. Signaling effectors underlying pathologic growth and remodeling of the heart. J Clin Invest. 2013;123:37–45.

5. Hill JA, Olson EN. Cardiac plasticity. N Engl J Med. 2008;358:1370–1380.

6. Akazawa H, Komuro I. Roles of cardiac transcription factors in cardiac hypertrophy. Circ Res. 2003;92:1079–1088.

7. Alexanian M, Padmanabhan A, McKinsey TA, Haldar SM. Epigenetic therapies in heart failure. J Mol Cell Cardiol. 2019;130:197–204.

8. Padmanabhan A, Haldar SM. Drugging transcription in heart failure. J Physiol [Internet]. 2019;Available from: http://dx.doi.org/10.1113/JP276745

9. Anand P, Brown JD, Lin CY, Qi J, Zhang R, Artero PC, Alaiti MA, Bullard J, Alazem K, Margulies KB, Cappola TP, Lemieux M, Plutzky J, Bradner JE, Haldar SM. BET bromodomains mediate transcriptional pause release in heart failure. Cell. 2013;154:569–582.

10. Spiltoir JI, Stratton MS, Cavasin MA, Demos-Davies K, Reid BG, Qi J, Bradner JE, McKinsey TA. BET acetyl-lysine binding proteins control pathological cardiac hypertrophy. J Mol Cell Cardiol. 2013;63:175–179.

11. Dhalluin C, Carlson JE, Zeng L, He C, Aggarwal AK, Zhou MM. Structure and ligand of a histone acetyltransferase bromodomain. Nature. 1999;399:491–496.

12. Taniguchi Y. The Bromodomain and Extra-Terminal Domain (BET) Family: Functional Anatomy of BET Paralogous Proteins. Int J Mol Sci [Internet]. 2016;17. Available from: http://dx.doi.org/10.3390/ijms17111849

13. Xu Y, Vakoc CR. Targeting Cancer Cells with BET Bromodomain Inhibitors. Cold Spring Harb Perspect Med [Internet]. 2017;7. Available from: http://dx.doi.org/10.1101/cshperspect.a026674

14. Stathis A, Bertoni F. BET Proteins as Targets for Anticancer Treatment. Cancer Discov. 2018;8:24–36.

15. Lovén J, Hoke HA, Lin CY, Lau A, Orlando DA, Vakoc CR, Bradner JE, Lee TI, Young RA. Selective inhibition of tumor oncogenes by disruption of super-enhancers. Cell. 2013;153:320–334.

16. Jang MK, Mochizuki K, Zhou M, Jeong H-S, Brady JN, Ozato K. The Bromodomain Protein Brd4 Is a Positive Regulatory Component of P-TEFb and Stimulates RNA Polymerase II-Dependent Transcription [Internet]. Molecular Cell. 2005;19:523–534. Available from: http://dx.doi.org/10.1016/j.molcel.2005.06.027

17. Yang Z, Yik JHN, Chen R, He N, Jang MK, Ozato K, Zhou Q. Recruitment of P-TEFb for Stimulation of Transcriptional Elongation by the Bromodomain Protein Brd4 [Internet]. Molecular Cell. 2005;19:535–545. Available from: http://dx.doi.org/10.1016/j.molcel.2005.06.029

18. Nicodeme E, Jeffrey KL, Schaefer U, Beinke S, Dewell S, Chung C-W, Chandwani R, Marazzi I, Wilson P, Coste H, White J, Kirilovsky J, Rice CM, Lora JM, Prinjha RK, Lee K, Tarakhovsky A. Suppression of inflammation by a synthetic histone mimic. Nature. 2010;468:1119–1123.

19. Filippakopoulos P, Qi J, Picaud S, Shen Y, Smith WB, Fedorov O, Morse EM, Keates T, Hickman TT, Felletar I, Philpott M, Munro S, McKeown MR, Wang Y, Christie AL, West N, Cameron MJ, Schwartz B, Heightman TD, La Thangue N, French CA, Wiest O, Kung AL, Knapp S, Bradner JE. Selective inhibition of BET bromodomains. Nature. 2010;468:1067–1073.

20. Duan Q, McMahon S, Anand P, Shah H, Thomas S, Salunga HT, Huang Y, Zhang R, Sahadevan A, Lemieux ME, Brown JD, Srivastava D, Bradner JE, McKinsey TA, Haldar SM. BET bromodomain inhibition suppresses innate inflammatory and profibrotic transcriptional networks in heart failure. Sci Transl Med [Internet]. 2017;9. Available from: http://dx.doi.org/10.1126/scitranslmed.aah5084

21. Antolic A, Wakimoto H, Jiao Z, Gorham JM, DePalma SR, Conner DA, Da Young L, Qi J, Seidman JG, Bradner JE, Brown JD, Haldar SM, Seidman CE, Burke MA. BET Bromodomain Proteins Regulate Transcriptional Reprogramming in Genetic Dilated Cardiomyopathy [Internet]. bioRxiv. 2020 [cited 2020 Apr 10];2020.02.09.940882. Available from: https://www.biorxiv.org/content/10.1101/2020.02.09.940882v1.abstract

22. Houzelstein D, Bullock SL, Lynch DE, Grigorieva EF, Wilson VA, Beddington RSP. Growth and early postimplantation defects in mice deficient for the bromodomain-containing protein Brd4. Mol Cell Biol. 2002;22:3794–3802.

23. Sohal DS, Nghiem M, Crackower MA, Witt SA, Kimball TR, Tymitz KM, Penninger JM, Molkentin JD. Temporally regulated and tissue-specific gene manipulations in the adult and embryonic heart using a tamoxifen-inducible Cre protein. Circ Res. 2001;89:20–25.

24. Koitabashi N, Bedja D, Zaiman AL, Pinto YM, Zhang M, Gabrielson KL, Takimoto E, Kass DA. Avoidance of transient cardiomyopathy in cardiomyocyte-targeted tamoxifen-induced MerCreMer gene deletion models. Circ Res. 2009;105:12–15.

25. Buenrostro JD, Giresi PG, Zaba LC, Chang HY, Greenleaf WJ. Transposition of native chromatin for fast and sensitive epigenomic profiling of open chromatin, DNA-binding proteins and nucleosome position. Nat Methods. 2013;10:1213–1218.

26. Huang B, Yang X-D, Zhou M-M, Ozato K, Chen L-F. Brd4 coactivates transcriptional activation of NF-kappaB via specific binding to acetylated RelA. Mol Cell Biol. 2009;29:1375–1387.

27. Shi J, Wang Y, Zeng L, Wu Y, Deng J, Zhang Q, Lin Y, Li J, Kang T, Tao M, Rusinova E, Zhang G, Wang C, Zhu H, Yao J, Zeng Y-X, Evers BM, Zhou M-M, Zhou BP. Disrupting the interaction of BRD4 with diacetylated Twist suppresses tumorigenesis in basal-like breast cancer. Cancer Cell. 2014;25:210–225.

28. Wu S-Y, Lee A-Y, Lai H-T, Zhang H, Chiang C-M. Phospho switch triggers Brd4 chromatin binding and activator recruitment for gene-specific targeting. Mol Cell. 2013;49:843–857.

29. Asangani IA, Dommeti VL, Wang X, Malik R, Cieslik M, Yang R, Escara-Wilke J, Wilder-Romans K, Dhanireddy S, Engelke C, Iyer MK, Jing X, Wu Y-M, Cao X, Qin ZS, Wang S, Feng FY, Chinnaiyan AM. Therapeutic targeting of BET bromodomain proteins in castration-resistant prostate cancer. Nature. 2014;510:278–282.

30. Roe J-S, Mercan F, Rivera K, Pappin DJ, Vakoc CR. BET Bromodomain Inhibition Suppresses the Function of Hematopoietic Transcription Factors in Acute Myeloid Leukemia. Mol Cell. 2015;58:1028–1039.

31. Heinz S, Benner C, Spann N, Bertolino E, Lin YC, Laslo P, Cheng JX, Murre C, Singh H, Glass CK. Simple combinations of lineage-determining transcription factors prime cis-regulatory elements required for macrophage and B cell identities. Mol Cell. 2010;38:576–589.

32. He A, Gu F, Hu Y, Ma Q, Ye LY, Akiyama JA, Visel A, Pennacchio LA, Pu WT. Dynamic GATA4 enhancers shape the chromatin landscape central to heart development and disease. Nat Commun. 2014;5:4907.

33. Akerberg BN, Gu F, VanDusen NJ, Zhang X, Dong R, Li K, Zhang B, Zhou B, Sethi I, Ma Q, Wasson L, Wen T, Liu J, Dong K, Conlon FL, Zhou J, Yuan G-C, Zhou P, Pu WT. A reference map of murine cardiac transcription factor chromatin occupancy identifies dynamic and conserved enhancers. Nat Commun. 2019;10:4907.

34. Arany Z, He H, Lin J, Hoyer K, Handschin C, Toka O, Ahmad F, Matsui T, Chin S, Wu P-H, Rybkin II, Shelton JM, Manieri M, Cinti S, Schoen FJ, Bassel-Duby R, Rosenzweig A, Ingwall JS, Spiegelman BM. Transcriptional coactivator PGC-1 alpha controls the energy state and contractile function of cardiac muscle. Cell Metab. 2005;1:259–271.

35. Lehman JJ, Boudina S, Banke NH, Sambandam N, Han X, Young DM, Leone TC, Gross RW, Lewandowski ED, Abel ED, Kelly DP. The transcriptional coactivator PGC-1alpha is essential for maximal and efficient cardiac mitochondrial fatty acid oxidation and lipid homeostasis. Am J Physiol Heart Circ Physiol. 2008;295:H185–96.

36. Takaya T, Kawamura T, Morimoto T, Ono K, Kita T, Shimatsu A, Hasegawa K. Identification of p300-targeted acetylated residues in GATA4 during hypertrophic responses in cardiac myocytes. J Biol Chem. 2008;283:9828–9835.

37. Lamonica JM, Deng W, Kadauke S, Campbell AE, Gamsjaeger R, Wang H, Cheng Y, Billin AN, Hardison RC, Mackay JP, Blobel GA. Bromodomain protein Brd3 associates with acetylated GATA1 to promote its chromatin occupancy at erythroid target genes. Proc Natl Acad Sci U S A. 2011;108:E159–68.

38. Gamsjaeger R, Webb SR, Lamonica JM, Billin A, Blobel GA, Mackay JP. Structural basis and specificity of acetylated transcription factor GATA1 recognition by BET family bromodomain protein Brd3. Mol Cell Biol. 2011;31:2632–2640.

39. Wu S-Y, Chiang C-M. The double bromodomain-containing chromatin adaptor Brd4 and transcriptional regulation. J Biol Chem. 2007;282:13141–13145.

40. Filippakopoulos P, Picaud S, Mangos M, Keates T, Lambert J-P, Barsyte-Lovejoy D, Felletar I, Volkmer R, Müller S, Pawson T, Gingras A-C, Arrowsmith CH, Knapp S. Histone recognition and large-scale structural analysis of the human bromodomain family. Cell. 2012;149:214–231.

41. Jung M, Philpott M, Müller S, Schulze J, Badock V, Eberspächer U, Moosmayer D, Bader B, Schmees N, Fernández-Montalván A, Haendler B. Affinity map of bromodomain protein 4 (BRD4) interactions with the histone H4 tail and the small molecule inhibitor JQ1. J Biol Chem. 2014;289:9304–9319.

42. Romero FA, Taylor AM, Crawford TD, Tsui V, Côté A, Magnuson S. Disrupting Acetyl-Lysine Recognition: Progress in the Development of Bromodomain Inhibitors. J Med Chem. 2016;59:1271–1298.

43. Dawson MA. The cancer epigenome: Concepts, challenges, and therapeutic opportunities. Science. 2017;355:1147–1152.

44. Piha-Paul SA, Sachdev JC, Barve M, LoRusso P, Szmulewitz R, Patel SP, Lara PN Jr, Chen X, Hu B, Freise KJ, Modi D, Sood A, Hutti JE, Wolff J, O’Neil BH. First-in-Human Study of Mivebresib (ABBV-075), an Oral Pan-Inhibitor of Bromodomain and Extra Terminal Proteins, in Patients with Relapsed/Refractory Solid Tumors. Clin Cancer Res. 2019;25:6309–6319.

45. Ray KK, Nicholls SJ, Buhr KA, Ginsberg HN, Johansson JO, Kalantar-Zadeh K, Kulikowski E, Toth PP, Wong N, Sweeney M, Schwartz GG, BETonMACE Investigators and Committees. Effect of Apabetalone Added to Standard Therapy on Major Adverse Cardiovascular Events in Patients With Recent Acute Coronary Syndrome and Type 2 Diabetes: A Randomized Clinical Trial. JAMA [Internet]. 2020;Available from: http://dx.doi.org/10.1001/jama.2020.3308

46. Winter GE, Buckley DL, Paulk J, Roberts JM, Souza A, Dhe-Paganon S, Bradner JE. DRUG DEVELOPMENT. Phthalimide conjugation as a strategy for in vivo target protein degradation. Science. 2015;348:1376–1381.

47. Lu J, Qian Y, Altieri M, Dong H, Wang J, Raina K, Hines J, Winkler JD, Crew AP, Coleman K, Crews CM. Hijacking the E3 Ubiquitin Ligase Cereblon to Efficiently Target BRD4. Chem Biol. 2015;22:755–763.

48. Zhou B, Hu J, Xu F, Chen Z, Bai L, Fernandez-Salas E, Lin M, Liu L, Yang C-Y, Zhao Y, McEachern D, Przybranowski S, Wen B, Sun D, Wang S. Discovery of a Small-Molecule Degrader of Bromodomain and Extra-Terminal (BET) Proteins with Picomolar Cellular Potencies and Capable of Achieving Tumor Regression. J Med Chem. 2018;61:462–481.

49. Stratton MS, Bagchi RA, Felisbino MB, Hirsch RA, Smith HE, Riching AS, Enyart BY, Koch KA, Cavasin MA, Alexanian M, Song K, Qi J, Lemieux ME, Srivastava D, Lam MPY, Haldar SM, Lin CY, McKinsey TA. Dynamic Chromatin Targeting of BRD4 Stimulates Cardiac Fibroblast Activation. Circ Res. 2019;125:662–677.

50. Garg V, Kathiriya IS, Barnes R, Schluterman MK, King IN, Butler CA, Rothrock CR, Eapen RS, Hirayama-Yamada K, Joo K, Matsuoka R, Cohen JC, Srivastava D. GATA4 mutations cause human congenital heart defects and reveal an interaction with TBX5. Nature. 2003;424:443–447.

51. Molkentin JD, Lin Q, Duncan SA, Olson EN. Requirement of the transcription factor GATA4 for heart tube formation and ventral morphogenesis. Genes Dev. 1997;11:1061–1072.

52. Ieda M, Fu J-D, Delgado-Olguin P, Vedantham V, Hayashi Y, Bruneau BG, Srivastava D. Direct reprogramming of fibroblasts into functional cardiomyocytes by defined factors. Cell. 2010;142:375–386.

53. Fu J-D, Stone NR, Liu L, Spencer CI, Qian L, Hayashi Y, Delgado-Olguin P, Ding S, Bruneau BG, Srivastava D. Direct reprogramming of human fibroblasts toward a cardiomyocyte-like state. Stem Cell Reports. 2013;1:235–247.

54. Qian L, Huang Y, Spencer CI, Foley A, Vedantham V, Liu L, Conway SJ, Fu J-D, Srivastava D. In vivo reprogramming of murine cardiac fibroblasts into induced cardiomyocytes. Nature. 2012;485:593–598.

55. Stone NR, Gifford CA, Thomas R, Pratt KJB, Samse-Knapp K, Mohamed TMA, Radzinsky EM, Schricker A, Ye L, Yu P, van Bemmel JG, Ivey KN, Pollard KS, Srivastava D. Context-Specific Transcription Factor Functions Regulate Epigenomic and Transcriptional Dynamics during Cardiac Reprogramming. Cell Stem Cell. 2019;25:87–102.e9.

56. Hashimoto H, Wang Z, Garry GA, Malladi VS, Botten GA, Ye W, Zhou H, Osterwalder M, Dickel DE, Visel A, Liu N, Bassel-Duby R, Olson EN. Cardiac Reprogramming Factors Synergistically Activate Genome-wide Cardiogenic Stage-Specific Enhancers. Cell Stem Cell. 2019;25:69–86.e5.

57. Oka T, Maillet M, Watt AJ, Schwartz RJ, Aronow BJ, Duncan SA, Molkentin JD. Cardiac-specific deletion of Gata4 reveals its requirement for hypertrophy, compensation, and myocyte viability. Circ Res. 2006;98:837–845.

58. Heineke J, Auger-Messier M, Xu J, Oka T, Sargent MA, York A, Klevitsky R, Vaikunth S, Duncan SA, Aronow BJ, Robbins J, Crombleholme TM, Molkentin JD. Cardiomyocyte GATA4 functions as a stress-responsive regulator of angiogenesis in the murine heart. J Clin Invest. 2007;117:3198–3210.

59. Neubauer S. The failing heart—an engine out of fuel. N Engl J Med. 2007;356:1140–1151.

60. Huss JM, Kelly DP. Mitochondrial energy metabolism in heart failure: a question of balance [Internet]. 2005;Available from: http://www.jci.org/articles/view/24405

61. Bates MGD, Bourke JP, Giordano C, d’Amati G, Turnbull DM, Taylor RW. Cardiac involvement in mitochondrial DNA disease: clinical spectrum, diagnosis, and management. Eur Heart J. 2012;33:3023–3033.

62. El-Hattab AW, Scaglia F. Mitochondrial Cardiomyopathies. Front Cardiovasc Med. 2016;3:25.

63. Dufour CR, Wilson BJ, Huss JM, Kelly DP, Alaynick WA, Downes M, Evans RM, Blanchette M, Giguère V. Genome-wide orchestration of cardiac functions by the orphan nuclear receptors ERRalpha and gamma. Cell Metab. 2007;5:345–356.

64. Stefanovic S, Christoffels VM. GATA-dependent transcriptional and epigenetic control of cardiac lineage specification and differentiation. Cell Mol Life Sci. 2015;72:3871–3881.

65. Shi J, Vakoc CR. The mechanisms behind the therapeutic activity of BET bromodomain inhibition. Mol Cell. 2014;54:728–736.

66. Faivre EJ, McDaniel KF, Albert DH, Mantena SR, Plotnik JP, Wilcox D, Zhang L, Bui MH, Sheppard GS, Wang L, Sehgal V, Lin X, Huang X, Lu X, Uziel T, Hessler P, Lam LT, Bellin RJ, Mehta G, Fidanze S, Pratt JK, Liu D, Hasvold LA, Sun C, Panchal SC, Nicolette JJ, Fossey SL, Park CH, Longenecker K, Bigelow L, Torrent M, Rosenberg SH, Kati WM, Shen Y. Selective inhibition of the BD2 bromodomain of BET proteins in prostate cancer. Nature. 2020;578:306–310.

67. Gilan O, Rioja I, Knezevic K, Bell MJ, Yeung MM, Harker NR, Lam EYN, Chung C-W, Bamborough P, Petretich M, Urh M, Atkinson SJ, Bassil AK, Roberts EJ, Vassiliadis D, Burr ML, Preston AGS, Wellaway C, Werner T, Gray JR, Michon A-M, Gobbetti T, Kumar V, Soden PE, Haynes A, Vappiani J, Tough DF, Taylor S, Dawson S-J, Bantscheff M, Lindon M, Drewes G, Demont EH, Daniels DL, Grandi P, Prinjha RK, Dawson MA. Selective targeting of BD1 and BD2 of the BET proteins in cancer and immuno-inflammation. Science [Internet]. 2020;Available from: http://dx.doi.org/10.1126/science.aaz8455

68. Shen C, Ipsaro JJ, Shi J, Milazzo JP, Wang E, Roe J-S, Suzuki Y, Pappin DJ, Joshua-Tor L, Vakoc CR. NSD3-Short Is an Adaptor Protein that Couples BRD4 to the CHD8 Chromatin Remodeler. Mol Cell. 2015;60:847–859.

69. Rahman S, Sowa ME, Ottinger M, Smith JA, Shi Y, Harper JW, Howley PM. The Brd4 extraterminal domain confers transcription activation independent of pTEFb by recruiting multiple proteins, including NSD3. Mol Cell Biol. 2011;31:2641–2652.

70. Lambert J-P, Picaud S, Fujisawa T, Hou H, Savitsky P, Uusküla-Reimand L, Gupta GD, Abdouni H, Lin Z-Y, Tucholska M, Knight JDR, Gonzalez-Badillo B, St-Denis N, Newman JA, Stucki M, Pelletier L, Bandeira N, Wilson MD, Filippakopoulos P, Gingras A-C. Interactome Rewiring Following Pharmacological Targeting of BET Bromodomains. Mol Cell. 2019;73:621–638.e17.

71. Love MI, Huber W, Anders S. Moderated estimation of fold change and dispersion for RNA-seq data with DESeq2. Genome Biol. 2014;15:550.

72. Chen EY, Tan CM, Kou Y, Duan Q, Wang Z, Meirelles GV, Clark NR, Ma’ayan A. Enrichr: interactive and collaborative HTML5 gene list enrichment analysis tool. BMC Bioinformatics. 2013;14:128.

73. Knowlton KU, Baracchini E, Ross RS, Harris AN, Henderson SA, Evans SM, Glembotski CC, Chien KR. Co-regulation of the atrial natriuretic factor and cardiac myosin light chain-2 genes during alpha-adrenergic stimulation of neonatal rat ventricular cells. Identification of cis sequences within an embryonic and a constitutive contractile protein gene which mediate inducible expression. J Biol Chem. 1991;266:7759–7768.

74. Burridge PW, Matsa E, Shukla P, Lin ZC, Churko JM, Ebert AD, Lan F, Diecke S, Huber B, Mordwinkin NM, Plews JR, Abilez OJ, Cui B, Gold JD, Wu JC. Chemically defined generation of human cardiomyocytes. Nat Methods. 2014;11:855–860.

